# Land-use intensity effects on the biodiversity-ecosystem functioning relationship in semi-natural grasslands at management-relevant spatial scales

**DOI:** 10.1101/2025.03.31.646360

**Authors:** Sophia N. Meyer, Javier Muro, Stephan Wöllauer, Lisa-Maricia Schwarz, Florian A. Männer, Olena Dubovyk, Anja Linstädter

## Abstract

**Context:** There is a limited understanding of how land-use intensity influences the relationship between plant diversity and the ecosystem function of aboveground biomass production in semi-natural grasslands at the field and farm scale. However, these spatial scales are critical to be addressed since management practices are applied at larger spatial scales than biophysical characteristics are measured.

**Objectives:** We aim to (1) examine the direct and indirect effects of land-use intensity on the spatial variability of biomass production, focusing on the role of plant species richness and its spatial variability at the field scale. We further aim to (2) compare the strength of the biodiversity-ecosystem functioning (BEF) relationship across spatial scales under different land-use intensities.

**Methods:** Going beyond the plot scale, we investigate the field and farm scale in two German regions for the years 2020 and 2021. Statistical analyses are based on prediction maps for species richness, biomass and land-use intensity, upscaled using Sentinel-2 satellite imagery, as well as on spectral dissimilarity (Rao’s Q) to account for environmental heterogeneity.

**Results:** High land-use intensity directly reduced species richness and its spatial variability, which acted as mediators, however, not as buffers for the spatial variability of biomass production at the field scale. The BEF relationship strengthened slightly with spatial scale at low land-use intensity, but is weakened under medium or even reversed under high intensity.

**Conclusions:** Our study provides new insights into how land-use intensity shapes the BEF relationship at spatial scales relevant for management, addressing spatial mismatches in complex social-ecological systems.

## Introduction

Semi-natural grasslands typically exhibit high biodiversity due to moderate and heterogeneous management practices (Bonari et al. 2017) and concurrently provide multiple ecosystem services (Le Provost et al. 2023; Prangel et al. 2024). As they are shaped by anthropogenic influence, these landscapes can be characterised as coupled social-ecological systems (SES) (Anderies et al. 2004), where ecological dynamics need to be understood in the context of social, economic and political aspects (Ostrom 2007). Ecological processes in grassland SES underlie a spatial mismatch between biophysical characteristics being measured at small spatial scales, and anthropogenic influence through management practices, operating at much larger scales (Cumming et al. 2006; Pelosi et al. 2010; Winkler et al. 2021; Klaus et al. 2024). Plant ecologists typically study ecological processes by sampling small plots, often only a few square meters in size and with limited spatial distribution (Abrahamson et al. 2011). However, generalising ecological processes from small to larger spatial scales can be misleading (Wheatley and Johnson 2009). Instead, scale-dependency must be explicitly considered by addressing the spatial scale at which environmental (Wiens 1989) and anthropogenic (Lindborg et al. 2017) drivers operate.

For the exploration of management practices in SES, a spatial upscaling (Fritsch et al. 2020) is required to match the field data as part of the ecological subsystem with the broader boundaries of the social subsystem (Martín-López et al. 2017; Klaus et al. 2024; Schulz et al. 2024). Land-use intensity in semi-natural grasslands is primarily determined by fertilisation (as input), as well as mowing and grazing (as output) (Blüthgen et al. 2012; Kuemmerle et al. 2013), all of which considerably influence plant diversity in semi-natural grasslands (Socher et al. 2012). Although the issue of spatial scale mismatch is increasingly recognized in the literature (Falco et al. 2022), the integration of knowledge from different disciplines poses a particular challenge to study ecological processes in grassland SES (Linstädter et al. 2016).

Management practices in European semi-natural grasslands are primarily regulated by policy, particularly through the Common Agricultural Policy of the European Union, which influences land use via agricultural subsidies and conservation measures (Vogt et al. 2019; Haensel et al. 2023). As a result, management practices are presumed to be more similar between fields belonging to the same farm than between fields belonging to different farms. Moreover, within individual grassland fields, management tends to be more homogeneous than between fields as management practices are adapted to site-specific characteristics such as slope (Andrieu et al. 2007). These two spatial scales, the field and farm scale, are therefore particularly relevant for research. Fields and farms can be regarded as management units, defined as spatially distinct areas that are managed as a coherent entity, usually for agricultural or conservation purposes (Klaus et al. 2024). However, the size of these units varies considerably across Europe, depending on the tenure system and climatic conditions of the respective SES (Kuemmerle et al. 2013).

Land-use intensification in Europe’s semi-natural grasslands is aimed at increasing biomass yields as an ecosystem service for society, but is also one of the main drivers of biodiversity loss (Wesche et al. 2012; Allan et al. 2014). At the plot scale, intensification typically homogenises grassland vegetation (Gossner et al. 2016) by threatening plant species with narrow ecological niches (Busch et al. 2019). In the long term, only a few competitive species persist, which reduces the plant diversity of the grassland (Wesche et al. 2012; Gerhold et al. 2013). The trade-off between biomass production and plant diversity in the context of land-use intensification is underpinned by the biodiversity-ecosystem functioning relationship (hereafter BEF relationship), or the diversity-productivity relationship (Tilman et al. 2014). Typically, a negative BEF relationship is observed under high land-use intensity and at small spatial scales, such as the plot scale (Beckmann et al. 2019; Andraczek et al. 2023). However, the direction and strength of the BEF relationship vary across spatial scales (Gross et al. 2000; Gonzalez et al. 2020; Thompson et al. 2021; Le Provost et al. 2023). Studies suggest that the BEF relationship tends to strengthen with spatial scale (Thompson et al. 2018; Craven et al. 2020), likely because the confounding effect of environmental heterogeneity diminishes at larger scales, allowing underlying ecological processes to be more robust (Wiens 1989; Stein et al. 2014). Yet, empirical investigations based on fields and farms as distinct spatial units remain scarce (but see Carmona et al. (2020)). The trade-off between biomass yield and plant species richness in response to mowing frequency was recently investigated at the scale of grassland fields (Schulz et al. 2024). However, spatial data on fields in this study originated from a segmentation approach using deep neural networks and remote sensing techniques (Tetteh et al. 2023) and therefore, do not correspond to the actual field boundaries. There remains a gap in understanding the impact of land-use intensity on the BEF relationship within the actual boundaries of fields and farms, likely due to the difficulty of obtaining spatial data on management units.

Land-use intensification leads to a decline in plant diversity, triggering a negative feedback loop decreasing ecosystem stability (Eisenhauer et al. 2024). In this context, plant diversity acts as biological insurance, as species respond differently to environmental changes, maintaining ecosystem functioning through functional redundancy (Wang and Loreau 2016; Loreau et al. 2021). A large species pool can therefore stabilise ecosystem functioning against increasing land-use intensity (Loreau et al. 2001), providing insurance both spatially (Loreau et al. 2021) and temporally, by enhancing resistance against environmental stressors (Caldeira et al. 2005; Hector et al. 2010) and facilitating dispersal (Daleo et al. 2023).

Ecological stability has numerous dimensions (Hillebrand et al. 2018). One measure, expressed as the inverse of variability, generally decreases from local to regional scales (Wang and Loreau 2014), though it remains unclear whether this pattern holds at intermediate spatial scales, such as at the field or farm scale. The covariance of species abundances, known as species synchrony, plays a major role in ecosystem stability (Walter et al. 2021). At small spatial scales, the spatial species synchrony of species richness has a stronger effect on the temporal stability of biomass production than species richness itself (Walter et al. 2021). Similarly, species covariation as the spatial equivalent of species synchrony (Daleo et al. 2023), has a stronger effect on the spatial variability of biomass production than species richness itself. From this perspective, a field or farm can be seen as a plant metacommunity composed of multiple local communities (Wang and Loreau 2014) which are linked through dispersal (Wilson 1992). Despite similar average species richness at the metacommunity level, differences in phylogenetic and functional composition can lead to varied species responses to disturbances such as mowing or grazing (Gerhold et al. 2013; Pouget et al. 2021). This suggests that understanding ecosystem stability at larger spatial scales requires accounting for the spatial variability of plant metacommunities.

This study examines the combined effects of land-use intensity on aboveground biomass production and plant species richness not only at the plot scale and its surroundings as done in Le Provost et al. (2023), but specifically at the field and farm scale and thus, on spatial scales that are particularly relevant for management in semi-natural grasslands. We use spatially upscaled biomass and plant species richness data (Muro et al. 2022), as well as upscaled land-use intensity data (Lange et al. 2022), generated by means of remote sensing and deep learning algorithms. Gonzalez et al. (2020) explicitly advocate the use of upscaled ecological data by means of remote sensing to analyse BEF relationships through space.

For this study, we adopt the concept of multi-grain alpha diversity (Sabatini et al. 2022), where field data are upscaled to pixels representing artificial sampling plots that function as virtual plant communities. As the species-area relationship and gamma diversity do not apply when grains are aggregated at larger spatial units (Sabatini et al. 2022), we introduce a new measure in our study, the spatial variability of species richness. This metric quantifies how niches are filled within a given spatial extent and how plant communities spatially respond to environmental fluctuations, independent of species identities. It is related to the spatial synchrony of species richness (Walter et al. 2021) and the spatial species covariation (Daleo et al. 2023), but is standardised and independent of the species-area relationship and thereby independent of size of the spatial unit. Finally, we build on the framework proposed by Klaus et al. (2024), which was developed for upscaling field-level data to the farm scale to inform agri-environmental policy. In contrast of aggregating only a limited number of a few field observations, we leverage a large multi-grain dataset to generate more robust predictions on how land-use intensity shapes BEF relationships in managed grasslands.

Our study aims to investigate both the direct and indirect effects of land-use intensity on species richness and the spatial variability of biomass production at the field scale, with a particular focus on the role of uniform management practices in semi-natural grasslands. Additionally, we address the spatial mismatch in social-ecological systems by analysing the BEF relationship at spatial scales most relevant for management decisions, specifically at the field and farm scale as well as at the commonly investigated plot scale. We hypothesise that:

(H1) Land-use intensity directly reduces plant species richness and its spatial variability at the field scale.
(H2) Plant species richness and its spatial variability within fields mediate the effects of land-use intensity on the spatial variability of biomass production.
(H3) The BEF relationship is stronger at larger spatial scales and under higher land-use intensity.

## Materials and methods

### Study area and design

The study area encompasses semi-natural grasslands of the biodiversity exploratories, a research platform situated in Central Europe consisting of three exploratory regions along a latitudinal gradient (Fischer et al. 2010). Our study focused on two exploratory regions, the biosphere reserve Schorfheide-Chorin (SCH) in north-eastern Germany and the Hainich-Dün region (HAI) in central Germany, situated within a national park (Fischer et al. 2010). Climate is temperate, with a mean annual temperature (MAT) in SCH ranging from 8-8.5 °C and a mean annual precipitation (MAP) of 500-600 mm. In contrast, HAI is slightly cooler with a MAT of 6.5-8 °C and a MAP of 500-800 mm (Fischer et al. 2010). While SCH is shaped by young glacial sediments with sandy and loamy soils, HAI is underlain by calcareous bedrock with loamy and clayey soils (Fischer et al. 2010). Grassland fields in both exploratories are distributed along a land-use intensity gradient (Blüthgen et al. 2012), ranging from extensively managed pastures to intensively used meadows. These fields are managed by local farmers, with land use determined by existing agricultural practices rather than experimental manipulations.

Our study focuses on three distinct spatial scales, the plot, field and farm scale. In each region there are 50 permanently installed grassland plots with a size of 50x50 m (Fischer et al. 2010). However, this study goes beyond fields containing only these plots and instead includes almost all fields and farms managed by local farmers in the regions SCH and HAI, which corresponds to an area of approximately 1,300 km^2^ (Fischer et al. 2010). To analyse processes at the plot scale with a sufficiently large sample size, we generated virtual plots of the same size as the permanently installed grassland plots (50x50 m) in almost every field. Similar to Schiller et al. (2024), we derived the boundaries of fields and farms from geospatial data of EU agricultural subsidy requests to the authorities of the German federal states Brandenburg (SCH) and Thuringia (HAI) for 2020 and 2021 respectively (Ministerium für Landwirtschaft, Umwelt und Klimaschutz des Landes Brandenburg (MLUK); Thüringer Landesamt für Landwirtschaft und Ländlichen Raum (TLLLR)). The spatial scales in this study follow a nested study design, where a virtual plot is located within in a field and several fields are associated with a respective farm (Fig. 1a). The spatial aggregation in virtual plots, fields and farms is based on upscaled data, which were generated by means of remote sensing and deep learning algorithms. Deep learning algorithms are a promising tool to predict vegetation data for whole regions (Cai et al. 2023; Dieste et al. 2024), or even to an entire country (Schulz et al. 2024), which could otherwise not be sampled in the field with this area coverage.

**Fig. 1.**
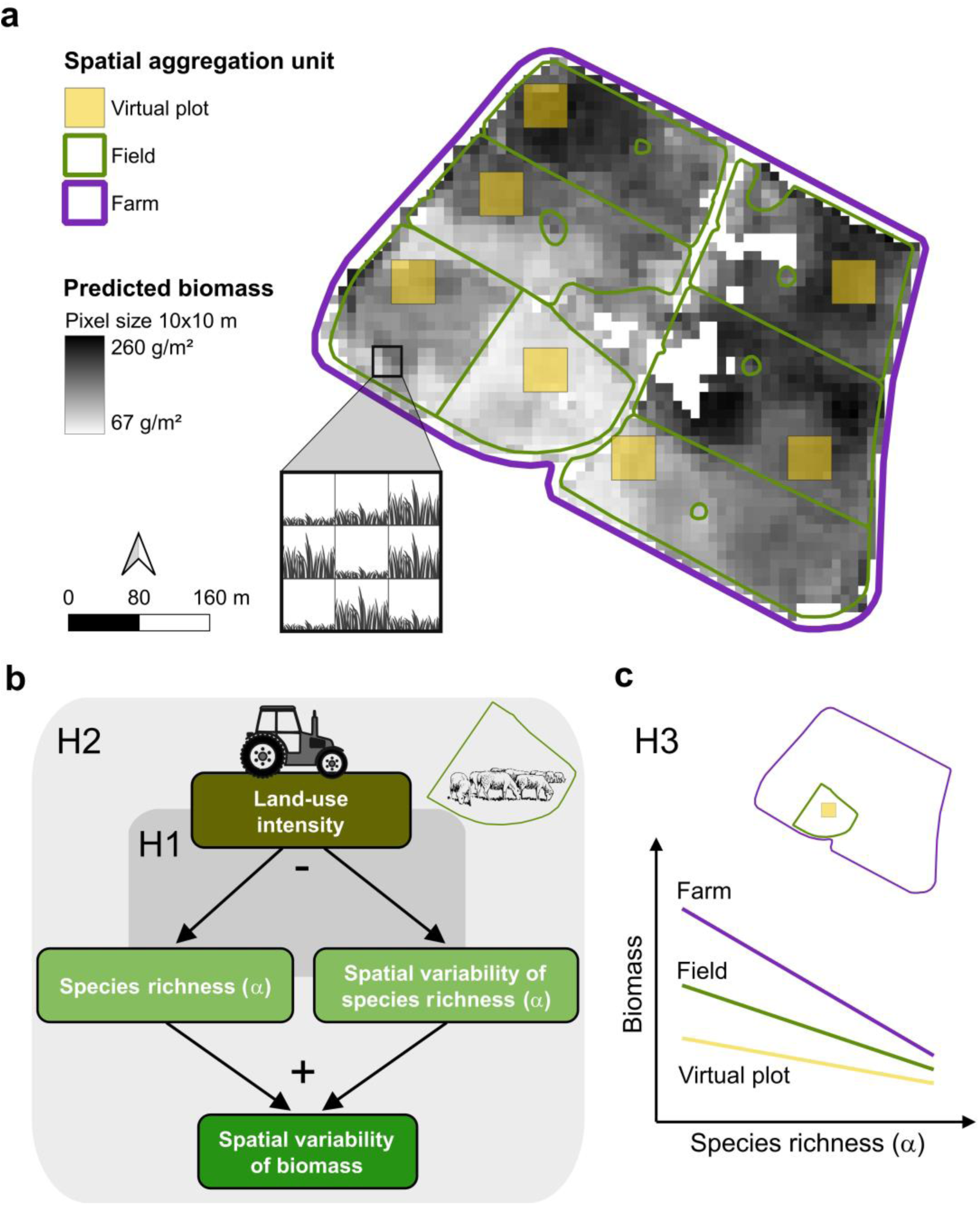
Conceptual framework for analysing the direct and indirect (biodiversity-mediated) effects of land-use intensity on the spatial variability of biomass production at the virtual plot, field and farm scale. (a) Prediction map of aboveground biomass as raster data (24-05-2020) in the exploratory region Schorfheide-Chorin and the three spatial aggregation scales. Pixels within the spatial units of virtual plots (yellow filled square), fields (green polygons) and farms (purple polygon) were aggregated to the mean or coefficient of variation (spatial variability). The 3x3 zoom window (black square) represents different biomass values. The illustrated farm is not representing the real association of fields due to data protection regulations. (b) Simplification of the conceptual structural equation model used to analyse direct and indirect land-use intensity effects referring to hypothesis H1 and H2. (c) Conceptual graph of comparing the relationship between plant species richness (alpha diversity) and biomass production across scales referring to hypothesis H3. Data source of biomass raster data: Muro et al. (2022). Data source of fields: Agricultural subsidy requests 2020 © MLUK, dl_de/by-2-0 (www.govdata.de/dl-de/by-2-0)/)

### Data acquisition

For the aggregation of pixel values to the respective spatial units, we first assembled upscaled data on aboveground biomass and species richness as well as on the land-use intensity index (LUI), spectral heterogeneity and topographic wetness index (TWI). All variables represent data of the years 2020 and 2021, except the LUI prediction map, which refers to the year 2018 (Table 1). In the following each variable used in this study is further described.

**Table 1.**
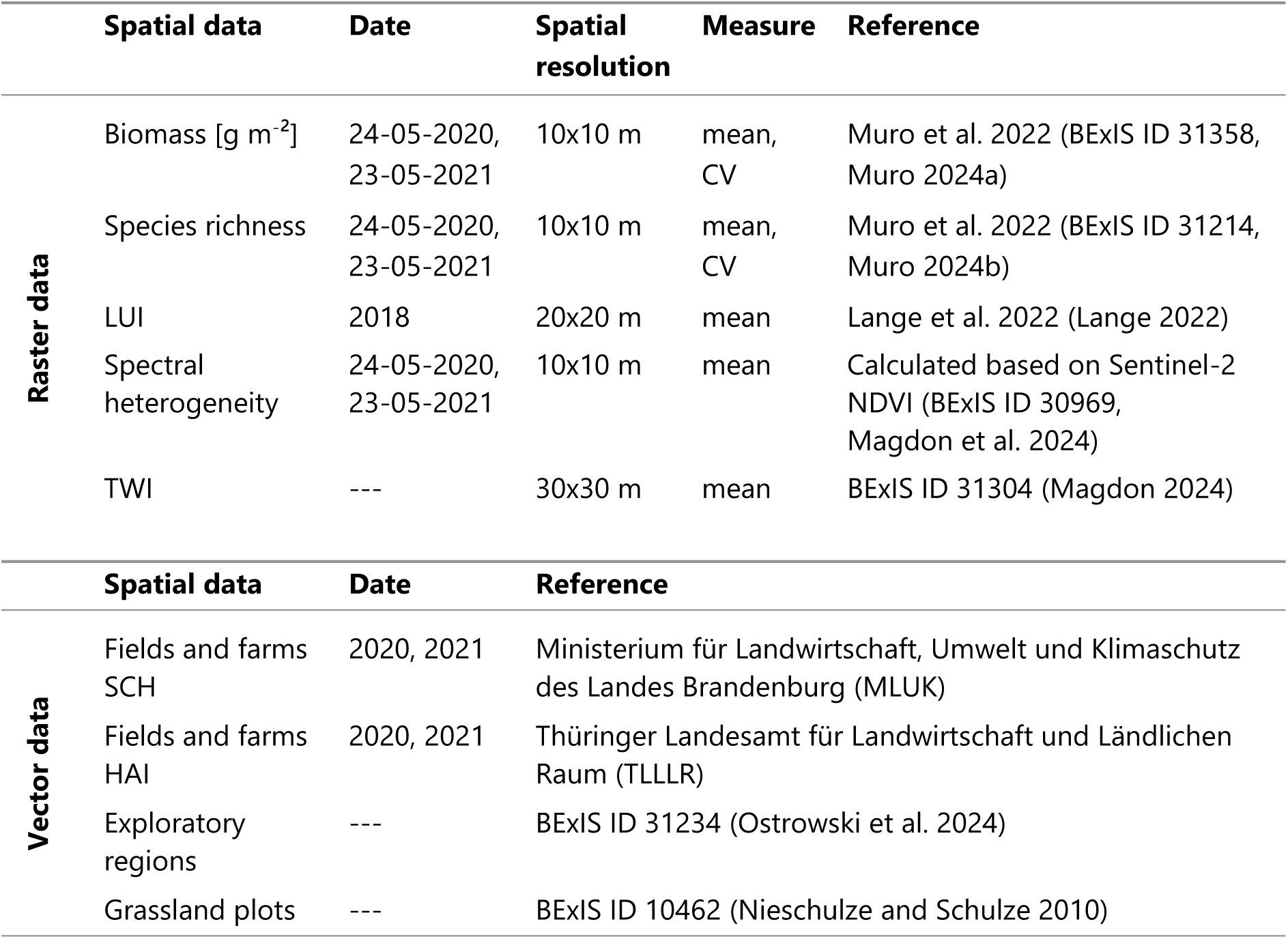
Set of upscaled data and vector data used for the spatial aggregation in this study. The mean or coefficient of variation (CV) is calculated for the raster datasets of biomass, species richness, land-use intensity index (LUI), spectral heterogeneity and topographic wetness index (TWI). Datasets with an ID are stored in the BExIS database (http://doi.org/10.17616/R32P9Q)

|  | Spatial data | Date | Spatial resolution | Measure | Reference |
| --- | --- | --- | --- | --- | --- |
| Raster data | Biomass [g m<sup>-2</sup>] | 24-05-2020, 23-05-2021 | 10x10 m | mean, CV | Muro et al. 2022 (BExIS ID 31358, Muro 2024a) |
|  | Species richness | 24-05-2020, 23-05-2021 | 10x10 m | mean, CV | Muro et al. 2022 (BExIS ID 31214, Muro 2024b) |
|  | LUI | 2018 | 20x20 m | mean | Lange et al. 2022 (Lange 2022) |
|  | Spectral heterogeneity | 24-05-2020, 23-05-2021 | 10x10 m | mean | Calculated based on Sentinel-2 NDVI (BExIS ID 30969, Magdon et al. 2024) |
|  | TWI | --- | 30x30 m | mean | BExIS ID 31304 (Magdon 2024) |
|  | Spatial data | Date | Reference |  |  |
| Vector data | Fields and farms SCH | 2020, 2021 | Ministerium für Landwirtschaft, Umwelt und Klimaschutz des Landes Brandenburg (MLUK) |  |  |
|  | Fields and farms HAI | 2020, 2021 | Thüringer Landesamt für Landwirtschaft und Ländlichen Raum (TLLLR) |  |  |
|  | Exploratory regions | --- | BExIS ID 31234 (Ostrowski et al. 2024) |  |  |
|  | Grassland plots | --- | BExIS ID 10462 (Nieschulze and Schulze 2010) |  |  |

#### Biomass and plant species richness

For the analysis of the BEF relationship at larger spatial scales we built on the upscaling approach of Muro et al. (2022). Aboveground biomass production and plant alpha diversity as species richness of 150 grassland plots was spatially upscaled with Sentinel-2 images (spatial resolution 10x10 m) by using a feed-forward neural network (see further details on upscaling method in Muro et al. (2022)). The upscaling was based on grassland vegetation records (Bolliger et al. 2021a), where plant species were inventoried in 16 m^2^ and biomass was destructively sampled in two subplots of 1 m^2^ (Bolliger et al. 2021b). The prediction model for biomass achieved an accuracy of R^2^ = 0.45 and the prediction on species richness achieved a model accuracy of R^2^ = 0.42 (further details in Muro et al. (2022)).

For our study, prediction maps of biomass were selected for the two time slices captured on 24 May 2020 and 23 May 2021, since the upscaled data originated from biomass cuttings conducted during the second half of May in both years (Muro et al. 2022). We did not summarise multiple time slices across the entire years of 2020 and 2021, as our primary focus was on within-site variation, which would not be accurately captured if data were aggregated over longer time periods. Pixel-wise biomass values from these prediction maps were aggregated into two proxies, the mean biomass [g m^-2^] (hereafter only biomass) and the coefficient of variation of biomass (hereafter biomass CV) at the virtual plot, field and farm scale. The mean of biomass was used to quantify biomass production in the respective spatial unit, while biomass CV served as the spatial variability of biomass production.

The upscaled plant species richness of Muro et al. (2022) represents multi-grain alpha diversity (Sabatini et al. 2022) since alpha diversity derived from vegetation surveys in 4x4 m built the basis for the upscaling (Bolliger et al. 2021a). The accumulation to gamma diversity cannot be implemented due to the species-area relationship, which implies a non-linear increase of alpha diversity with area (Sabatini et al. 2022). The mean of species richness (hereafter richness) provides the information how many species co-occur on average in a given area. Our newly introduced measure, the spatial variability of species richness (hereafter richness CV), is used as a comparable proxy for the relative variability of alpha diversity in differently sized management units. Similar to the spectral variation hypothesis (Palmer et al. 2002), which assumes a positive relationship between the dissimilarity across pixels and species diversity, richness CV is used in our study as an indicator of how diverse ecological niches are filled within an area. The CV of species richness was also regarded in previous studies as dominance index of a plant community (Thukral et al. 2019). However, it must be noted that richness CV cannot be directly compared with beta diversity or species covariation (Daleo et al. 2023) since information of species identities is missing.

#### Land-use intensity

Upscaled data on land-use intensity were obtained from Lange et al. (2022). These data were based on yearly interviews on land use with the owners of fields on which grassland plots of the biodiversity exploratories are located. The interview data build the basis for the calculation of the land-use intensity index (LUI) (Blüthgen et al. 2012; Vogt et al. 2019; Ostrowski et al. 2020). This continuous index is the sum of the standardised components mowing, grazing and fertilisation (Blüthgen et al. 2012). The study of Lange et al. (2022) spatially upscaled all the three land-use intensity components to the whole of Germany by means of convolutional neural networks and Sentinel-2 images for the years 2017 and 2018. They calculated the overall LUI based on those three prediction maps and achieved an overall prediction accuracy of R^2^ = 0.82 (Lange et al. 2022). For the analysis of this study, the upscaled LUI of the year 2018 was used (Lange et al. 2022). Aggregated means of LUI above five were regarded as outliers and removed from the dataset before analysis.

#### Spectral heterogeneity

We quantified spectral heterogeneity (Rocchini et al. 2010) as a proxy for spatial environmental heterogeneity (Torresani et al. 2024). It was calculated as Rao’s quadratic entropy (Rao’s Q) (Rocchini et al. 2017) based on the normalised vegetation index (NDVI) of preprocessed Sentinel-2 satellite images (Magdon et al. 2024):

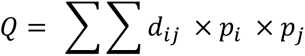

For that, the same time slices of Sentinel-2 images as for biomass and species richness were selected to match with the biotic variables and because spatial heterogeneity of grassland productivity in spring was found to be especially high (Klaus et al. 2016). The NDVI raster layers with 32-bit resolution were transformed to 8-bit resolution (values from 0-255). Subsequently, the dissimilarity index Rao’s Q was calculated as Euclidean distance (pairwise distance (d_ij_) between relative abundance of pixel values i and j) with the R package rasterdiv (Rocchini et al. 2021) based on a moving window of 9x9. This measure is based on the pure spectral signal of vegetation, whereas our variable biomass CV was upscaled based on collected field data. Thus, spectral heterogeneity is not treated as biodiversity indicator and thereby does not refer to the spectral variation hypothesis (Palmer et al. 2002) as mentioned above. According to Fahrig et al. (2011), spectral heterogeneity can be considered as compositional heterogeneity. This measure was included in the analysis to control for the fact that species diversity is influenced by the availability of niches provided by a habitat (Stein et al. 2014), referring to Hutchinson’s concept of the niche (Hutchinson 1957). We calculated mean values of spectral heterogeneity across all three spatial scales.

#### Topographic wetness index

We used the topographic wetness index (TWI) as a proxy for soil moisture (Raduła et al. 2018), as it integrates the effects of topography and soil characteristics which are decisive factors in shaping plant communities (Araya et al. 2011; Oddershede et al. 2015). At larger spatial scales, such as at farm scale, terrain characteristics are known to play a more important role than soil characteristics (Zhu and Lin 2011), making the TWI meaningful for analysing larger spatial scales. TWI was derived from digital elevation models and calculated by Magdon (2024) for the whole exploratory regions (see further details in the metadata of Magdon (2024)). In our study, we calculated the mean of the TWI to control for environmental conditions in the statistical models.

#### Spatial aggregation

We extracted pixels from the upscaled data for each plot, field and farm polygon corresponding to the three spatial scales (Fig. 1a). These pixel values were then aggregated to calculate the mean and standard deviation, from which we derived the coefficient of variation (CV) for selected variables. At first, we filtered fields classified as meadows, pastures, mown pastures and other grasslands. To ensure representative aggregation values for species richness and biomass, we filtered fields with at least 80% of pixel coverage within the polygons. Anonymised identification numbers of farmers’ agricultural subsidy requests provided information on field ownership, allowing us to group fields into their respective farms (MLUK; TLLLR). The composition of fields within a given farm and the grassland type vary between years. We randomly created virtual plots (50x50 m), which represent the smallest spatial scale. However, the area of virtual plots (2500 m^2^) is larger than the area of the subplots (16 m^2^) in which the vegetation surveys for the upscaling were done (Bolliger et al. 2021a; Muro et al. 2022). Virtual plots were generated using QGIS v3.22 (QGIS.org 2022) by placing a square centred on a random point at least 50 m from field boundaries. Due to the small size of some fields, not every field contained a virtual plot (see Table S1).

Each polygon layer was transformed to the coordinate reference system of the respective raster layer before pixel extraction. We applied the summing or averaging approach according to Klaus et al. (2024) to acquire zonal statistics, though we computed means and not area-weighted means as suggested. Pixel extraction and aggregation were performed using the raster R package (Hijmans 2023).

### Data analysis

All statistical analyses were conducted in R (R Core Team 2022). For the analyses, the dataset was split into four subsets based on region and year, accounting for regional differences and data dependency across years.

#### Direct and indirect effects of land-use intensity

To evaluate the direct and indirect effects of land-use intensity on biomass CV at the field scale, we tested piecewise structural equation models (pSEM), a specific type of path analysis, using the piecewiseSEM R package (Lefcheck 2016). We formulated a conceptual model of hypothesised causal relationships based on a-priori knowledge (Table S3), where LUI, spectral heterogeneity, field size and TWI were included as exogenous variables. Richness, richness CV and biomass were included as mediators, whereas biomass CV represented the overall response variable (Fig. 1b). We fitted four separate pSEMs for each region and year, since observations within the same region are not completely independent due to similar environmental characteristics between years (Lefcheck 2016). Removing data dependence by using mixed effect models with only two factor levels as random effect is proven to inaccurately calculate the estimates (Harrison 2015) and was therefore not used for the analysis. The initial four models based on our a-priori conceptual model (Fig. S1) were not acceptable. Therefore, for each model additional paths with potential ecological relevance were added until they reached a Fisher’s C statistic with a p-value above 0.05 (Shipley 2009). To fulfill the model assumptions on normal distribution and homoscedasticity of residuals for each multiple regression in the respective pSEMs, some variables needed to be square root or logarithmically transformed.

#### BEF relationship across spatial scales

We performed multiple linear regressions separately for each region and year to test the relationship between richness and biomass as well as between richness and biomass CV across spatial scales and at different land-use intensities (Fig. 1c). We tested a three-way interaction between richness, spatial scale and LUI to allow a direct comparison of the slope of the BEF relationship. Prior to analysis, the continuous LUI variable was classified into the three categories low (0-1.82), medium (1.82-3.27) and high (3.27-4.98) LUI for a better interpretability of the interaction. The variables spectral heterogeneity, size of the spatial unit and TWI were included as control. Here again, a linear mixed effect model with a hierarchical structure could not be implemented due to few factor levels, despite the nested structure. To meet the model assumptions of linearity and homoscedasticity of residuals, some variables were accordingly square root or logarithmically transformed.

To validate whether the slope of the BEF relationships is similar between upscaled data and data measured in the field, the grassland plots (50x50 m) of the biodiversity exploratories were examined. Pixels were extracted and aggregated in the plot boundaries and compared to field measurements of the respective grassland plot by using multiple linear regressions (more details on methodology in Online Resource).

## Results

We found that both richness CV and biomass CV increased from virtual plots over fields to farms in both regions (Fig. S4). Biomass and richness did not significantly differ between the regions at the respective spatial scales (Fig. S4). At the field and farm scale, the region HAI was generally higher regarding LUI and spectral heterogeneity compared to SCH (Fig. S4). We detected that SCH had up to 1.39 ha larger fields and 8.88 ha larger farms compared to HAI (Table S2, Fig. S4), accompanied by a higher number of fields and farms in HAI (Table S1). Regarding the grassland type of fields, there were relatively more pastures and mown pastures in HAI and less meadows compared to SCH (Table S1).

### Direct and indirect effects of land-use intensity

To distinguish between the direct and indirect effects of LUI at the field scale, we tested four pSEMs, analysing different subsets based on region and year. All four final models were found to have good model fit statistics (see Table S4 for a direct comparison of pSEMs and Table S5-8 for more details). We found a direct negative effect of LUI on richness across all subsets with standardised path coefficients (β) ranging from -0.15 in SCH 2021 to -0.29 in HAI 2020. We also observed a direct negative effect of LUI on richness CV in all subsets except for HAI 2021. These effects were rather weak with β ranging from -0.06 in HAI 2020 to -0.13 in SCH 2021. Since pSEMs had slightly different paths and were tested based on different subsets, the most parsimonious model cannot be identified according to Akaike’s information criterion (AIC). Similarly, the effect sizes allow only a limited comparison between models. Among the models, HAI 2020 (Fig. 2b) explained the highest proportion of variance for richness (R^2^ = 0.19) and richness CV (R^2^ = 0.34), whereas it explained the least variance for biomass CV (R^2^ = 0.29). The highest explained variance for biomass CV was detected in the SCH 2020 model (R^2^ = 0.52) (Fig. 2a).

**Fig. 2.**
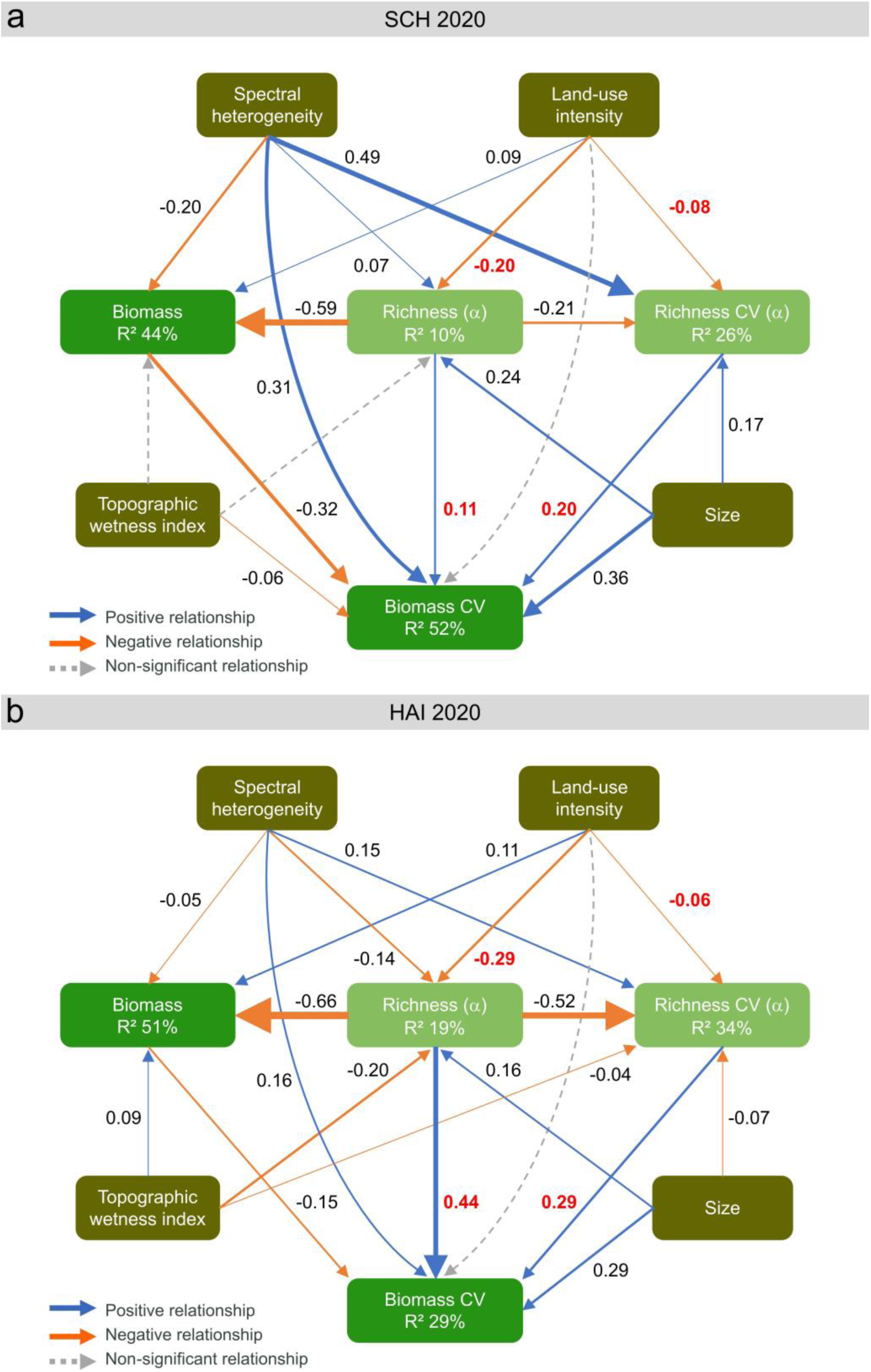
Piecewise structural equation models (pSEMs) for the analysis of direct and indirect land-use intensity effects on the spatial variability of biomass (biomass CV). pSEMs were separately tested for each subset based on the regions (a) Schorfheide-Chorin (SCH) and (b) Hainich-Dün (HAI) and the year 2020. Model fit statistics of pSEM for the subset SCH 2020: Fisher’s C = 9.19, AIC (d-sep) = 11,665.72, df = 6, p-value = 0.163, N = 1,304. Model fit statistics of pSEM for subset HAI 2020: Fisher’s C = 10.526, AIC (d-sep) = 9,361.32, df = 6, p-value = 0.104, N = 1,969. The thickness of paths corresponds to the magnitude of standardised estimates, which are illustrated next to path arrows. Mediator variables are displayed in the second row of the pSEMs. Estimates showing the magnitude of paths addressed in the first and second hypothesis are coloured in red. Note that standardised estimates cannot be directly compared in terms of units since variables were transformed differently (see Table S5 and Table S7), models tested different data subsets and the structure of pSEMs was slightly different

We detected mediation effects of both richness and richness CV for the indirect effect of LUI on biomass CV, though they were relatively weak and statistically significant in only three of the four models. Indirect effects were calculated as the product of the standardised path coefficients, and total mediation effects represent the sum of indirect paths involving the mediator variables (Lefcheck 2016). The first component of this indirect effect corresponds to the negative direct impact of LUI on richness and richness CV. The second component is the direct effect of richness or richness CV on biomass CV, which was unexpectedly positive in all models. The strongest direct effect regarding the mediator richness was found in HAI 2020 (β = 0.44), whereas no effect was found for HAI 2021 (Fig. S2). The mediator richness CV had a direct effect on biomass CV with intermediate effect sizes ranging from 0.20 in SCH 2020 to 0.29 in HAI 2020. In three of the four models, the direct positive effect of richness CV on biomass CV was even stronger than that of species richness itself.

The indirect negative effects of LUI on biomass CV, mediated by richness, were detected in all models except in HAI 2021. Although these effects were overall weak, the magnitude in HAI 2020 (β = -0,13) was about ten times stronger than in the other two significant models (SCH 2020: β = -0.02; SCH 2021: β = -0.01). Similarly, weak indirect effects of LUI on biomass CV, mediated by richness CV, were observed (SCH 2020: β = -0.02; SCH 2021: β = -0.04; HAI 2020: β = -0.02). The total mediation effect of both richness variables regarding the indirect effect of LUI on biomass CV was highest in HAI 2020 (β = -0.15), compared to SCH 2020 (β = -0.04) and SCH 2021 (β = -0.03). The direct effect of LUI on biomass CV cannot be compared to the indirect effects mediated by the two richness variables since this path was not significant (p > 0.05) in all models.

Regarding the other control variables, spectral heterogeneity had a positive direct effect on richness CV in all models, with the strongest effect in SCH 2020 (β = 0.49). In contrast, the relationship between spectral heterogeneity and richness was inconsistent and generally weak across models. Furthermore, spectral heterogeneity showed a small to intermediate direct negative effect on biomass CV in all models, with the strongest effect again in SCH 2020 (β = 0.31). Pearson’s correlation analyses revealed that correlations between spectral heterogeneity and both the richness and biomass variables did not exceed 0.3 (Fig. S3).

### BEF relationship across spatial scales

Consistent with the findings from pSEMs, we observed negative relationships between biomass and species richness across all regression models, except for SCH in 2021 at the farm scale and under high LUI (Fig. 3, model fit statistics in Table S9-12). We only detected differences between slopes of the BEF relationship across spatial scales and LUI for the SCH models at high LUI. For SCH in 2021, we found that the slope of the BEF relationship was slightly steeper at the virtual plot scale than at the field scale (Table S10), which contradicts our expectation of steeper slopes at larger spatial scales. This significant three-way interaction detected in SCH 2021 means that under high LUI, a unit increase in square root transformed biomass corresponds to a decrease of 0.52 units in square root transformed species richness at the field scale, whereas at the virtual plot scale, the decrease was notably higher by 2.56 units.

**Fig. 3.**
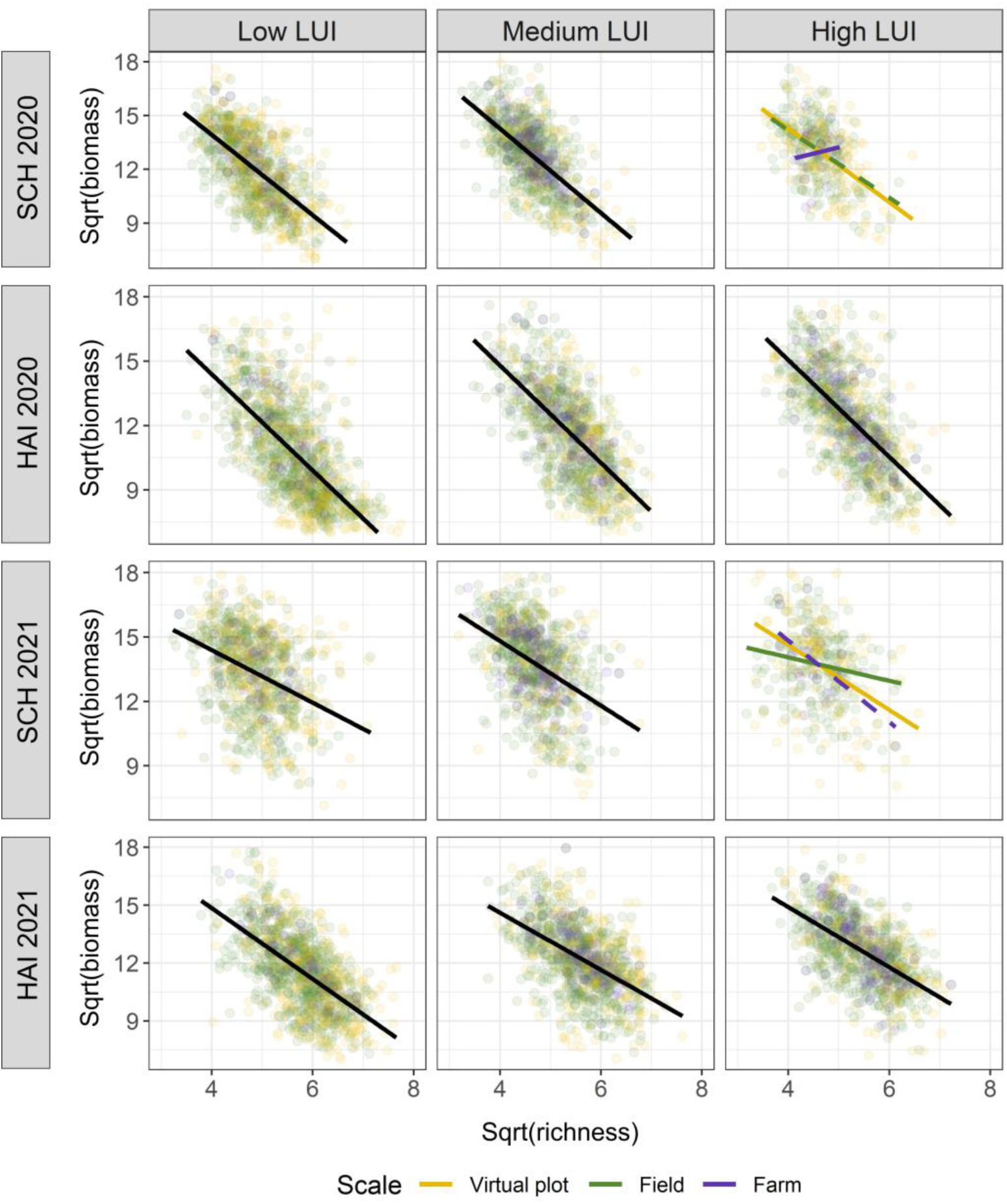
Relationships between biomass and species richness across the virtual plot (yellow), field (green) and farm (purple) scale at different land-use intensities (LUI). Separate multiple linear regressions were conducted for each subset based on the regions Schorfheide-Chorin (SCH) and Hainich-Dün (HAI) in 2020 and 2021, testing a three-way interaction between species richness, spatial scale and LUI (SCH 2020: R^2^ = 0.46***, F20,1983 = 87.43; HAI 2020: R^2^ = 0.51***, F20,2861 = 148.7; SCH 2021: R^2^ = 0.19***, F20,1967 = 23.66; HAI 2021: R^2^ = 0.39***, F20,2842 = 92.82). Points display raw data of bivariate relationships, while lines show fitted values from regression models. Black lines indicate significant relationships between biomass and richness. Coloured and dashed regression lines indicate a significant interaction between richness and scale, whereas only coloured solid lines indicate a significant three-way interaction

We found varying slopes regarding the BEF relationship between richness and biomass CV across spatial scales for all models except for SCH 2021 (Fig. 4, Table S13-16). The strength of the BEF relationship slightly increased with spatial scale, but was unexpectedly positive, similar to what was found in pSEMs. The three-way interaction between richness, spatial scale and LUI was only significant for SCH 2020 at medium LUI (Table S13) and for HAI 2021 at high LUI (Table S16). In SCH 2020, the steepest slope was observed at the farm scale, though it was weaker at medium compared to low LUI. The model of HAI 2021 in contrast, had a steeper slope at the virtual plot scale under high LUI compared to low LUI. Under high LUI, this model exhibited an increase of 0.22 units square root transformed biomass CV per unit square root transformed species richness at the field scale, whereas a higher increase of 0.42 units at the virtual plot scale. Regarding low LUI, this model showed a weak increase of 0.10 units at virtual plot scale versus a much higher increase of 0.46 units at the field scale.

**Fig. 4.**
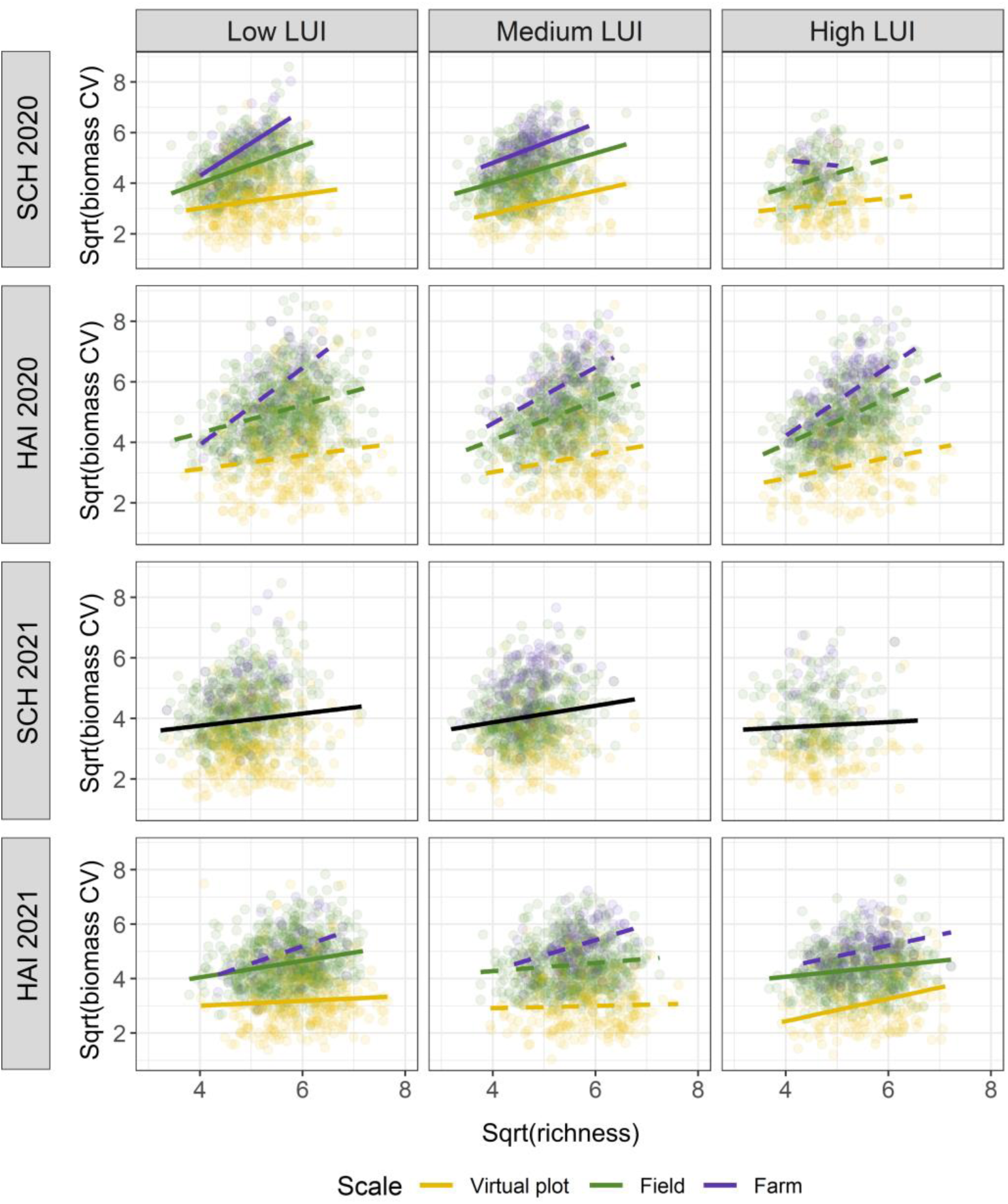
Relationship between spatial variability of biomass (biomass CV) and species richness across the virtual plot (yellow), field (green), and farm (purple) scale at different land-use intensities (LUI). Separate multiple linear regressions were conducted for each subset based on the regions Schorfheide-Chorin (SCH) and Hainich-Dün (HAI) in 2020 and 2021, testing a three-way interaction between species richness, spatial scale and LUI (SCH 2020: R^2^ = 0.58***, F20,1983 = 139.60; HAI 2020: R^2^ = 0.44***, F20,2861 = 116.10; SCH 2021: R^2^ = 0.43***, F20,1967 = 76.99; HAI 2021: R^2^ = 0.45***, F20,2842 = 116.30). Points display raw data of bivariate relationships, while lines show fitted values from regression models. Black lines indicate significant relationships between biomass and richness. Coloured and dashed regression lines indicate a significant interaction between richness and scale, whereas only coloured solid lines indicate a significant three-way interaction

We detected a consistent negative correlation between biomass and spectral heterogeneity, whereas a positive correlation between biomass CV and spectral heterogeneity, similar to the trend of the respective richness variables. Comparing standardised effect sizes across all eight regression models, richness had the strongest influence on biomass, while spatial scale had the strongest effect on biomass CV (Table S17). All other variables, including the interaction between spatial scale and richness, had very small effect sizes (<0.3).

The validation of BEF relationships in the permanent grassland plots were significant for the models which were based on the upscaled data, whereas not significant when they were based on measured field data (Fig. S5-6, Table S18-19). Thus, a direct comparison of the BEF relationship between upscaled and measured data was not possible.

## Discussion

In our study, we take scale-dependent effects of land-use intensity on the biodiversity-ecosystem functioning (BEF) relationship into account by specifically addressing the field and farm scale, which are focal scales for understanding and generalising ecological processes (Wheatley and Johnson 2009) in semi-natural grasslands. We found that land-use intensity directly decreased species richness and its spatial variability, which in turn positively affected the spatial variability of biomass production. The comparison of the BEF relationship revealed that scale-dependent patterns are especially pronounced under low land-use intensity, with the strongest relationship at the farm scale.

### Direct effects of land-use intensity

In line with our first hypothesis, land-use intensity had a direct negative effect on species richness and its spatial variability at the field scale. This finding is supported by Carmona et al. (2020), who specifically investigated field-level intensity in a European arable system and reported that plant species richness decreased threefold from the lowest to the highest land-use intensity. The negative impact of land-use intensity on species richness was slightly stronger in the Hainich-Dün region compared to Schorfheide-Chorin, likely due to undetected intensification effects in the land-use intensity index of the region Schorfheide-Chorin. This discrepancy may be explained by substantial nutrient releases from historical wetland drainage without the application of fertilisers, in contrast to more intense fertilising in Hainich-Dün (Lamers et al. 2002; Blüthgen et al. 2012). Furthermore, management practices differ between the two regions, with a higher mean grazing intensity in Schorfheide-Chorin and a higher mean mowing frequency in Hainich-Dün (Vogt et al. 2019). Nevertheless, we found a consistent, moderately strong negative effect across years and regions, providing robust evidence for biotic homogenisation of multi-grain alpha diversity at the field scale. This aligns with studies reporting a spatial homogenisation of plant diversity in response to land-use intensification (Gossner et al. 2016; Liu et al. 2018; Busch et al. 2019), typically addressing the plot scale (Wesche et al. 2012; Beckmann et al. 2019). Habitat heterogeneity counteracts the negative effects of land-use intensity on species richness (Maskell et al. 2019). However, in our study, spectral heterogeneity as a proxy for environmental heterogeneity was surprisingly negatively correlated with species richness in three of the four path models. Negative relationships between spectral variation and species richness have been reported for larger spatial extents (Schmidtlein and Fassnacht 2017; Fassnacht et al. 2022), which may explain this finding.

While our proposed measure of the spatial variability of species richness was not as strongly affected by land-use intensity as species richness itself, it still confirmed a biotic homogenisation of grassland communities within fields. Blüthgen et al. (2016) documented a negative relationship between land-use intensity and temporal species asynchrony in bats and birds, albeit not in plants. Spatial species synchrony, which is more similar to our proposed measure of the spatial variability of species richness, may be more important than the more frequently studied temporal species synchrony when addressing larger spatial scales.

### Direct diversity-stability relationships

Contrary to our expectations, species richness and its spatial variability increased the spatial variability of biomass production, instead of decreasing it. This contrasts with the findings of Daleo et al. (2023), who demonstrated that alpha and gamma diversity reduced the spatial variability of biomass production in grasslands, whereas beta diversity increased it. Conversely, Qiao et al. (2022) found a positive relationship between alpha diversity and stability for larger spatial scales in temperate forests, albeit they did not calculate the coefficient of variation (CV). Our proposed measure of the spatial variability of species richness exhibited the same positive relationship as reported for beta diversity (Daleo et al. 2023). Its similarity to beta diversity may explain the observed pattern, however, it does not incorporate species identities and thus cannot detect dissimilarities in species composition. A simulation study showed that both alpha and beta diversity may exert a spatial insurance effect, buffering for ecosystem variability at regional scales (Wang and Loreau 2016). In the context of the BEF relationship, beta diversity is more strongly associated with biomass than alpha diversity at increasing spatial scales due to environmental filtering (Reu et al. 2022; Liang et al. 2022; Daleo et al. 2023). This could explain our finding of stronger direct relationships between the spatial variability of biomass and the spatial variability of species richness within fields since this measure is more similar to beta diversity.

The positive relationship between the spatial variability of biomass and both species richness variables may be affected by underlying environmental heterogeneity, distorting upscaled pixel values (Markham et al. 2023). Consistent with our results, remote sensing studies observed that the NDVI as a productivity index was negatively correlated with species richness, whereas its CV as a measure for spatial heterogeneity, was positively correlated (Oindo and Skidmore 2002). The positive relationship detected between spectral heterogeneity as a proxy for environmental heterogeneity and the spatial variability of biomass does not necessarily mean that spatial ecosystem stability decreases. Furthermore, there was a relatively strong positive effect of spectral heterogeneity on the spatial variability of species richness, especially in the region Schorfheide-Chorin. A reason for this could be that the field size is on average greater in Schorfheide-Chorin, as also shown for fields in entire Brandenburg by Jänicke et al. (2024). Increasing the extent of aggregation reduces the variation of a variable (Hobbs et al. 2007) which thus, can better reveal patterns in larger than in smaller patches. Moreover, a statistical averaging effect (or portfolio effect) is considered to be present, which implies a generally positive diversity-stability relationship (CV as a reversed measure for stability) due to the aggregation of biomass at community level (Tilman et al. 1998; Doak et al. 1998) and was assumed to be the main effect in shaping the diversity-stability relationship (Zhao et al. 2022). However, this mathematical relationship is refuted to be exclusively positive (Lhomme and Winkel 2002) and was so far rather addressed in relation to the temporal stability of ecosystems.

### Indirect effects of land-use intensity

We can confirm our second hypothesis that plant species richness and its spatial variability mediate the effects of land-use intensity on the spatial variability of biomass. Unexpectedly, we found an overall negative indirect effect of land-use intensity, which increased spatial biomass stability. However, we could not detect an indirect buffering effect of species richness and its spatial variability as the direct land-use intensity effect on the spatial variability of biomass was not significant and thus, could not be compared to the indirect effect. The indirect effect was generally weak, but strongest in the Hainich-Dün region in 2020. This finding may be explained by increased spatial heterogeneity in Hainich-Dün due to a higher proportion of pastures on which cattle graze (Pouget et al. 2021). Contrary to our expectations, the results show that the two richness variables do not directly reduce, but increase the spatial variability of biomass production. This positive direct effect, which is part of the indirect effect, can be attributed to a hidden effect of spatial heterogeneity that is captured in remotely sensed species richness.

The reasons for the detected weak indirect effect of land-use intensity on the spatial variability of biomass are three-fold. First, a selection bias may be present since site characteristics already select for certain land use practices in respect of how intense a grassland can be used (Andrieu et al. 2007; Blüthgen et al. 2012). For instance, patches naturally high in soil fertility may be more suitable for intensified management. In contrast, patches of steep slope are more suitable for extensive management, e.g. by sheep grazing (Fischer et al. 2010), and are therefore more spatially heterogeneous (Nunes et al. 2019), ultimately increasing the spatial variability of biomass. Second, intensive management practices can lead to a homogenisation effect of soil and vegetation properties within fields (Pouget et al. 2021), decreasing the availability of niches. Hence, it does not mean that species diversity is not buffering the spatial variability of biomass, but that it is overlayed by the homogenisation effect at high land-use intensities. Structural equation models showed that spectral heterogeneity as a proxy for spatial environmental heterogeneity strongly promoted the spatial variability of biomass, acting as an antagonist of land-use intensity (see Gossner et al. (2016) who reported similar effects for plant diversity). Third, the upscaled LUI data only represents a single year (2018) and are not congruent for the upscaled biomass and diversity data of 2020 and 2021. As land-use intensity can vary significantly between years (Vogt et al. 2019), a temporal mismatch between land-use intensity data and ecological data might have occurred. Moreover, a single year may not fully capture long-term management effects. Hence, an upscaled land-use intensity index averaged over a longer period might better reflect vegetation responses.

### BEF relationship across spatial scales

Consistent with our third hypothesis and findings of Thompson et al. (2018) and Craven et al. (2020), we found that the commonly reported negative BEF relationship between biomass and species richness (Andraczek et al. 2023) strengthened with spatial scale. However, this trend was only evident under low land-use intensity and the effect was rather weak. At medium and at high land-use intensity, the relationship between biomass and species richness was unexpectedly stronger at the virtual plot scale compared to the field scale, though only evident for the region Schorfheide-Chorin. This means that the factor of high LUI attenuates the effect of spatial scale increasing the strength of the BEF relationship. This pattern partly aligns with Gross et al. (2000), who aggregated sampling plots to the field scale and provided evidence for a negative linear BEF relationship with species diversity expressed as species densities. However, when increasing the scale to whole biogeographic regions, they detected unimodal relationships (Gross et al. 2000). This highlights that aggregation scale and sample size matter because increasing the grain size decreases the variance in an area (Wiens 1989), ultimately revealing stronger relationships when differences between areas become higher. However, the extent of variance reduction with increasing grain size depends on the spatial heterogeneity of the area (Wiens 1989). Moreover, it is difficult to separate heterogeneity effects from the size of a unit due to the generally positive area-heterogeneity relationship (Stein et al. 2014). At high land-use intensity, species richness declines due to reduced heterogeneity (Meynard et al. 2013), which was confirmed by our study’s strongest negative BEF relationship under low land-use intensity at the farm scale.

The positive relationship between species richness and the spatial variability of biomass, which can be considered as the opposite of stability, was also stronger under low land-use intensity and at the field and farm scale. Although the relationship was reversed, Liang et al. (2022) reported similar findings that the buffering effect of plant diversity on the community stability increases from local to regional scales. In contrast, the average ecosystem stability is reported to be highest at the local scale when diversity decreases at larger spatial scales (Wang and Loreau 2014), e.g. as a consequence of high land-use intensity and thereby less heterogeneous environments. As our study represents aggregated multi-grain alpha diversity (Sabatini et al. 2022) and not gamma diversity, species richness can be reduced at larger spatial scales. This trend could explain our finding that the relationship between species richness and the spatial variability of biomass weakened with increasing spatial scale under high land-use intensity.

### Limitations and outlook

Our study has several methodological limitations. Most of our variables were derived from Sentinel-2 satellite imagery with coarse pixel sizes of 10x10 m, thus certain dependencies may be incorporated in our statistical models. As this study is based on predicted data, some unexplained variance remains, with about 50% of variance being unexplained in the upscaling approach of species richness and biomass (Muro et al. 2022). For instance, spatial heterogeneity is an inherent part of the upscaling and cannot be avoided when using remote sensing techniques (Markham et al. 2023). Although the large sample size in our study can provide greater statistical power, biases in data due to spatial heterogeneity or remote sensing limitations might still persist. It could be useful to acknowledge that while a larger sample size improves statistical confidence, it does not completely resolve issues related to the accuracy of predictions. Discussing how the results might differ with different sample sizes or data sources could be a way to critically reflect on this compensation. Furthermore, for future studies we recommend to calculate area-weighted means of upscaled data within spatial units to better reflect patch specific characteristics, as suggested in Klaus et al. (2024).

Scale dependency remains a challenge when environmental and anthropogenic drivers operate at different spatial scales than biotic responses are measured (Wiens 1989; Lindborg et al. 2017). Aggregating upscaled data to the field and farm scale can help mitigate this issue, as presented in our study. Further ecological studies addressing the field and farm scale are meaningful, particularly regarding the BEF relationship and the role of plant functional diversity in the spatial and temporal stability of ecosystem functioning (de Bello et al. 2021).

## Conclusion

Upscaled data representing metacommunities in form of multi-grain alpha diversity (Sabatini et al. 2022) display a great potential to address the BEF relationship as well as ecosystem stability at spatial scales of management interventions. Our study provides evidence for the homogenisation effect of land-use intensity on multi-grain alpha diversity at the field scale. For this scale, we identified mediation effects of species richness and the newly proposed measure of the spatial variability of species richness on the spatial variability of biomass production, however, no indirect buffering effect was found. Furthermore, our results show that the BEF relationship is stronger at the field and farm scale compared to the virtual plot scale under low land-use intensity, revealing scale-dependent effects of land-use intensity.

This study highlights the demand for further research conducting empirical analyses at management-relevant spatial scales with the overarching goal to provide recommendations for agri-environmental policy, for instance in the context of regulating agricultural subsidies. From a practical perspective, our study approach could lay the basis for reinforcing ‘spatially targeted policy instruments’ at the regional scale, as proposed by Huber et al. (2022). Such policy instruments could help to compensate for certain proportions of less productive grasslands, or to modify landscapes by creating incentives for environmental heterogeneity, e.g. through the subdivision of large fields by different mowing events as proposed by Bonari et al. (2017). Greater incentives for environmental heterogeneity in terms of economic aspects could be more firmly anchored in the European Union’s Common Agricultural Policy as a prerequisite to maintain and promote plant diversity, which in turn warrants the spatial stability of biomass production that is crucial for farmers.

## Acknowledgements

We thank the managers of the three Exploratories, Julia Bass, Max Müller, Anna K. Franke, Miriam Teuscher, Robert Künast, Franca Marian, Uta Schumacher, Melissa Jüds and all former managers for their work in maintaining the plot and project infrastructure; Victoria Grießmeier for giving support through the central office, Andreas Ostrowski for managing the central data base, and Markus Fischer, Eduard Linsenmair, Dominik Hessenmöller, Daniel Prati, Ingo Schöning, François Buscot, Ernst-Detlef Schulze, Wolfgang W. Weisser and the late Elisabeth Kalko for their role in setting up the Biodiversity Exploratories project. We thank the administration of the Hainich national park, the UNESCO Biosphere Reserve Swabian Alb and the UNESCO Biosphere Reserve Schorfheide-Chorin as well as all land owners for the excellent collaboration. The work has been funded by the DFG Priority Program 1374 ‘Biodiversity-Exploratories’ (grant code LI 1842/4–1, DU 1596/1–1 and MA 6917/2-5). Field work permits were issued by the responsible state environmental offices of Baden-Württemberg, Thüringen, and Brandenburg.

We thank the Botany core project for providing vegetation data for the upscaling of plant species richness and biomass as performed in Muro et al. (2022). We thank Juliane Helm for her helpful feedback on an earlier version of the manuscript. Special thanks to the authorities MLUK (Ministerium für Landwirtschaft, Umwelt und Klimaschutz des Landes Brandenburg) and TLLLR (Thüringer Landesamt für Landwirtschaft und Ländlichen Raum) for providing geospatial data on agricultural field and farm boundaries from agricultural subsidy requests.

## Statements and declarations

### Author contributions

Writing of the first draft of the manuscript: SNM. Conceptualisation of the study: SNM, AL, OD. Data preparation and data analysis: SNM. Data collection of biomass and species richness as part of project Sensing Biodiversity Across Scales (SeBAS): SNM, JM, LS, FM, AL. SW provided data on processed Sentinel-2 images and maps on topographic wetness index. All authors commented on and edited previous versions of the manuscript.

### Data availability

This work is based on data elaborated by the project Sensing Biodiversity Across Scales (SeBAS) and the project Instrumentation and Remote Sensing of the Biodiversity Exploratories program (DFG Priority Program 1374). The datasets 31234 (polygons of borders of exploratory regions) and 31304 (topographic wetness index) are publicly available in the Biodiversity Exploratories Information System (http://doi.org/10.17616/R32P9Q). However, to give data owners and collectors time to perform their analysis the Biodiversity Exploratories’ data and publication policy includes by default an embargo period of three years from the end of data collection/data assembly which applies to the datasets: 10462 (polygons of grassland plots of biodiversity exploratories), 31214 (predicted plant species richness 2017-2021), 31358 (predicted grassland biomass 2020-2021), 30969 (pre-processed Sentinel-2 satellite images), 31891 (aggregated pixel values at virtual plot, field and farm scale) and 31895 (aggregated field data of biomass and species richness from field surveys and pixel values at plot scale). These datasets will be made publicly available via the same data repository.

Polygons of grassland fields and farms originate from geospatial data of agricultural subsidy requests provided by the authorities of Brandenburg (Ministerium für Landwirtschaft, Umwelt und Klimaschutz des Landes Brandenburg (MLUK), Henning-von-Tresckow-Straße 2-13,14467 Potsdam) and Thuringia (Thüringer Landesamt für Landwirtschaft und Ländlichen Raum (TLLLR), Naumburger Str. 98, 07743 Jena) for the years 2020 and 2021. Polygons for grassland fields in Brandenburg are publicly available at https://geobroker.geobasis-bb.de/gbss.php?MODE=GetProductInformation&PRODUCTID=996f8fd1-c662-4975-b680-3b611fcb5d1f. The authors do not have permission to share the data of farm polygons for Brandenburg as well as the field and farm polygons for Thuringia.

Prediction maps on land-use intensity raster data were published in Lange et al. (2022) and is publicly available at the UFZ repository, https://ufz.maps.arcgis.com/apps/webappviewer/index.html?id=192195ae64534ff9ae655082b6145774.

Figures were created by Sophia N. Meyer with Inkscape (https://inkscape.org/), QGIS v3.22 (QGIS.org 2022) and the R package ggplot2 (Wickham H (2016) Elegant Graphics for Data Analysis. Springer-Verlag New York. https://ggplot2.tidyverse.org/).

