## Supplementary file for "Land-use intensity effects on the biodiversity-ecosystem functioning relationship in semi-natural grasslands at management-relevant spatial scales"

**Title:**

**Journal:**

Landscape Ecology

**Authors:**

Sophia N. Meyer[1*], Javier Muro[2,3], Stephan Wöllauer[4], Lisa-Maricia Schwarz[1,5], Florian A. Männer[1,5,6], Olena Dubovyk[2,7], Anja Linstädter[1]

**Affiliations:**

[1] Biodiversity Research/ Systematic Botany, University of Potsdam, Maulbeerallee 1, 14469 Potsdam, Germany

[2] Center for Remote Sensing of Land Surfaces (ZFL), University of Bonn, Genscherallee 3, 53113 Bonn, Germany

[3] Thünen-Institute of Farm Economics, Bundesallee 63, 38116 Braunschweig, Germany

[4] Faculty of Resource Management, HAWK University of Applied Sciences and Arts, Büsgenweg 1a, 37077 Goettingen, Germany

[5] Institute of Crop Science and Resource Conservation, University of Bonn, Karlrobert-Kreiten-Str. 13, 53115 Bonn, Germany

[6] Fraunhofer Institute for Computer Graphics Research IGD, Competence Center Bioeconomy- Smart Farming, Joachim-Jungius-Str. 11, 18059 Rostock, Germany

[7] Terrestrial Remote Sensing, University of Hamburg, Bundesstr. 55, 20146 Hamburg, Germany

***Corresponding author:**

Sophia N. Meyer,

Biodiversity Research/ Systematic Botany, University of Potsdam, Maulbeerallee 1, 14469 Potsdam, Germany

**Supplementary methods**

*R code of function for automated pixel extraction and aggregation*

```
PixelExtraction <- function(raster, polygon) {
 # Detect cores and create a cluster
 cores <- detectCores()
 clust <- makeCluster(cores - 1)

 # Dynamically capture the names of the raster and polygon objects
 raster_name <- deparse(substitute(raster))
 polygon_name <- deparse(substitute(polygon))
 clusterExport(clust, c(raster_name, polygon_name))

 # Transform sf dataset to SpatialLayer
 polygon <- as_Spatial(polygon)

 # Prepare the results data frame
 stats_name <- paste0("stats_", deparse(substitute(raster)))
 stats <- as.data.frame(polygon)
 assign(stats_name,stats, envir = .GlobalEnv) #envir = .GlobalEnv necessary to define since
function works in new environment

 # Perform parallel extraction and statistics calculation
 results <- parSapply(clust, 1:length(polygon), function(i) {
   extracted <- raster::extract(raster, polygon[i,])

   extracted_num <- as.numeric(unlist(extracted))
   mean = mean(extracted_num, na.rm = TRUE)
   sum = sum(extracted_num, na.rm = TRUE)
   sd = sd(extracted_num, na.rm = TRUE)
   CV = sd / mean * 100
   list(mean = mean, sum = sum, sd = sd, CV = CV)
 }, USE.NAMES = TRUE)

 # Stop the cluster
 stopCluster(clust)

 # Format the results
 results_long <- as.data.frame(results)
 results_long <- t(results_long)
 results_long <- as.data.frame(results_long)

 # Retrieve the dynamically named stats data frame
 stats_df <- get(stats_name)

 # Bind the stats data frame to the results_long data frame
 combined_df <- cbind(stats_df, results_long)
```

```
# Unnest list in data frame
combined_df <- combined_df %>% unnest(cols = c(tail(names(combined_df), 4)))

return(combined_df)
}
```

*Details on model selection of piecewise structural equation models (pSEMs)*

Meaningful paths that were not considered in the a-priori conceptual model were added by checking the independence claim with the lowest p-value and identifying the missing path compared to the tested pSEM. This procedure was repeated until d-separation of the basis set was fulfilled (Shipley 2009). Paths were added to the a-priori conceptual pSEM in the following order. First, for all models, the path *richness ~ size* needed to be added. Secondly, for the two SCH models the path *biomass CV ~ TWI* was included, whereas for the two HAI models the path *richness CV ~ TWI* was added. A third additional path (*richness CV ~ biomass*) needed to be added for the subset HAI 2021, which led to a smaller number of degrees of freedom compared to the other three models.

*Predicted and measured BEF relationship in grassland plots of the biodiversity exploratories*

The slope of the BEF relationships between species richness and biomass as well as between species richness and biomass CV was compared between predicted data and actually measured data in grassland plots of the biodiversity exploratories. For the model based on predicted data, pixels were extracted in the plot polygons and aggregated to the mean and coefficient of variation (CV). For the model based on measured data, measurements from the third exploratory region Schwäbische Alb (ALB) were also included to obtain a higher number of field observations. Destructive biomass was measured in three squares (0.6x0.6 m) per plot, for which the mean for 1 $m^2$ was calculated to retrieve biomass values per plot. Field measurements on biomass CV originated from Rising Plate Meter (RPM) measurements (unit represents 0.5 cm increments) in ten squares (1 $m^2$) along a 50 m transect of the plot. Non-destructive RPM measurements in the range from 0-60 were filtered before they were transformed to biomass estimations by using the calibration equation 'Estimated biomass = 3.11 + 1.06 * RPM' (Table S18-19, Fig. S5-6). Predicted species richness represents an area of 16 $m^2$, whereas the measured species richness represents an area of 3 $m^2$. Biomass is expressed in g $m^{-2}$ for both predicted and measured data. For each response variable (biomass or biomass CV) and data type (predicted or measured data), separate multiple linear regressions were analysed which included the year and region as control variables.

### Supplementary figures

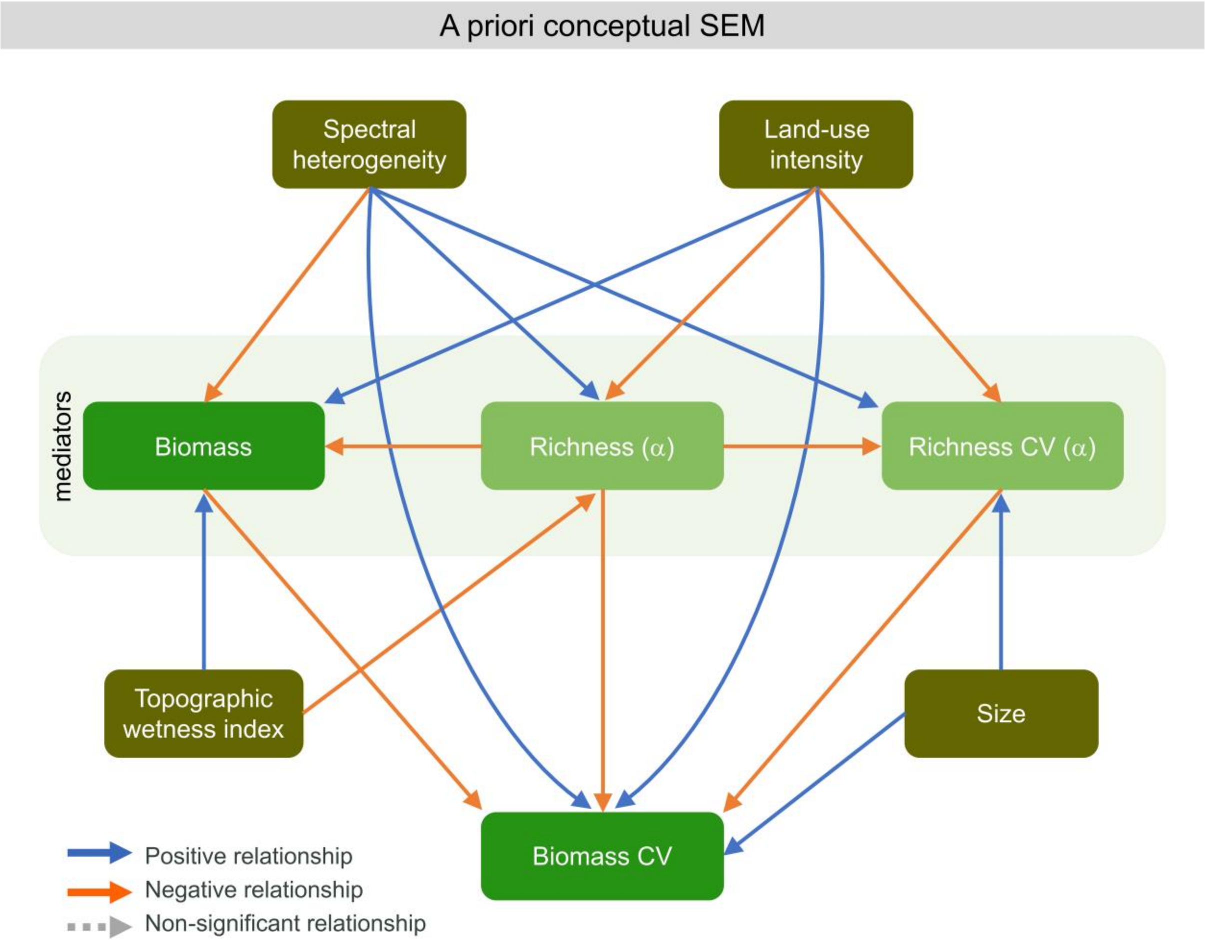


**Fig. S1** A-priori conceptual structural equation model (SEM) for the analysis of direct and indirect land-use effects on the spatial variability of biomass production (biomass CV). The mediator variables species richness and its spatial variability (richness CV) as well as biomass are displayed in the second row of the pSEMs. Paths are based on a literature review (see Table S3)

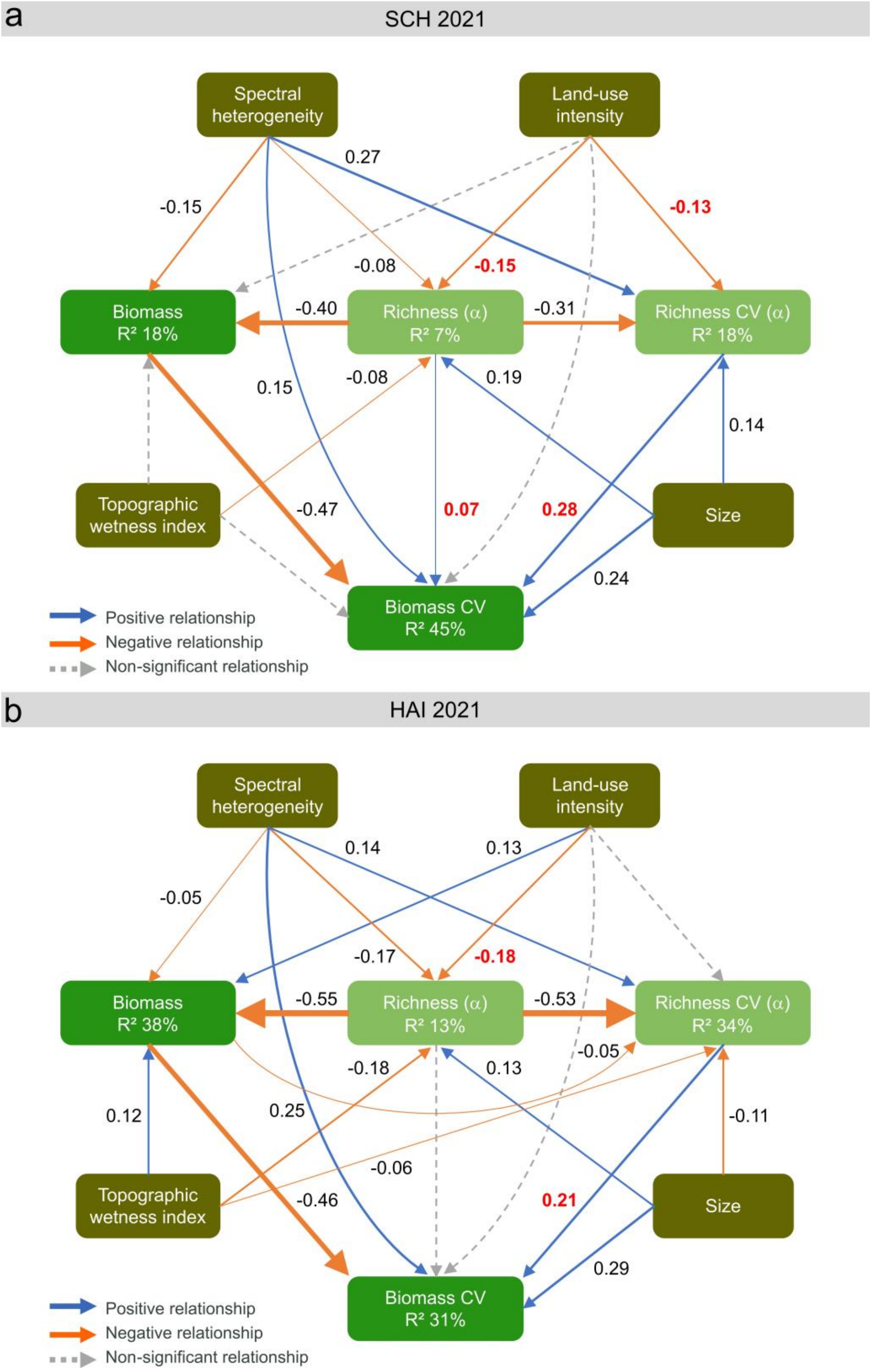


**Fig. S2** Piecewise structural equation models (pSEM) for the analysis of direct and indirect land-use intensity effects on the spatial variability of biomass (biomass CV). pSEMs were separately tested for each subset based on the regions (a) Schorfheide-Chorin (SCH) and (b) Hainich-Dün (HAI) and the year 2021. Model fit statistics of pSEM for the subset SCH 2021: Fisher's C =10.559, AIC (d-sep) = 19,257.43, df = 6, p-value = 0.103, N = 1,283. Model fit statistics of pSEM for subset HAI 2021: Fisher's C = 6.502, AIC (d-sep) = 8,627.08, df = 4, p-value = 0.165, N = 1,961. The thickness of paths corresponds to the magnitude of standardised estimates, which are illustrated next to path arrows. Mediator variables are displayed in the second row of the pSEMs. Estimates showing the magnitude of paths addressed in the first and second hypothesis are coloured in red. Note that standardised estimates cannot be directly compared in terms of units since variables were transformed differently (see Table S6 and Table S8), models tested different data subsets and the structure of pSEMs was slightly different

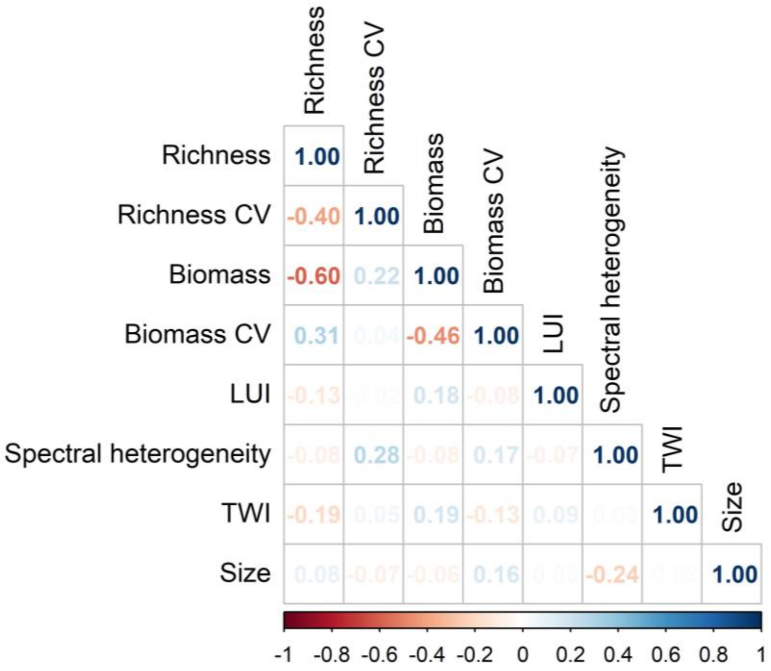


**Fig. S3** Pierson's correlation coefficients of all variables included in piecewise structural equation models for testing direct and indirect land-use intensity effects on the spatial variability of biomass at the field scale. Correlations were calculated based on the whole dataset and were not separated by year and region

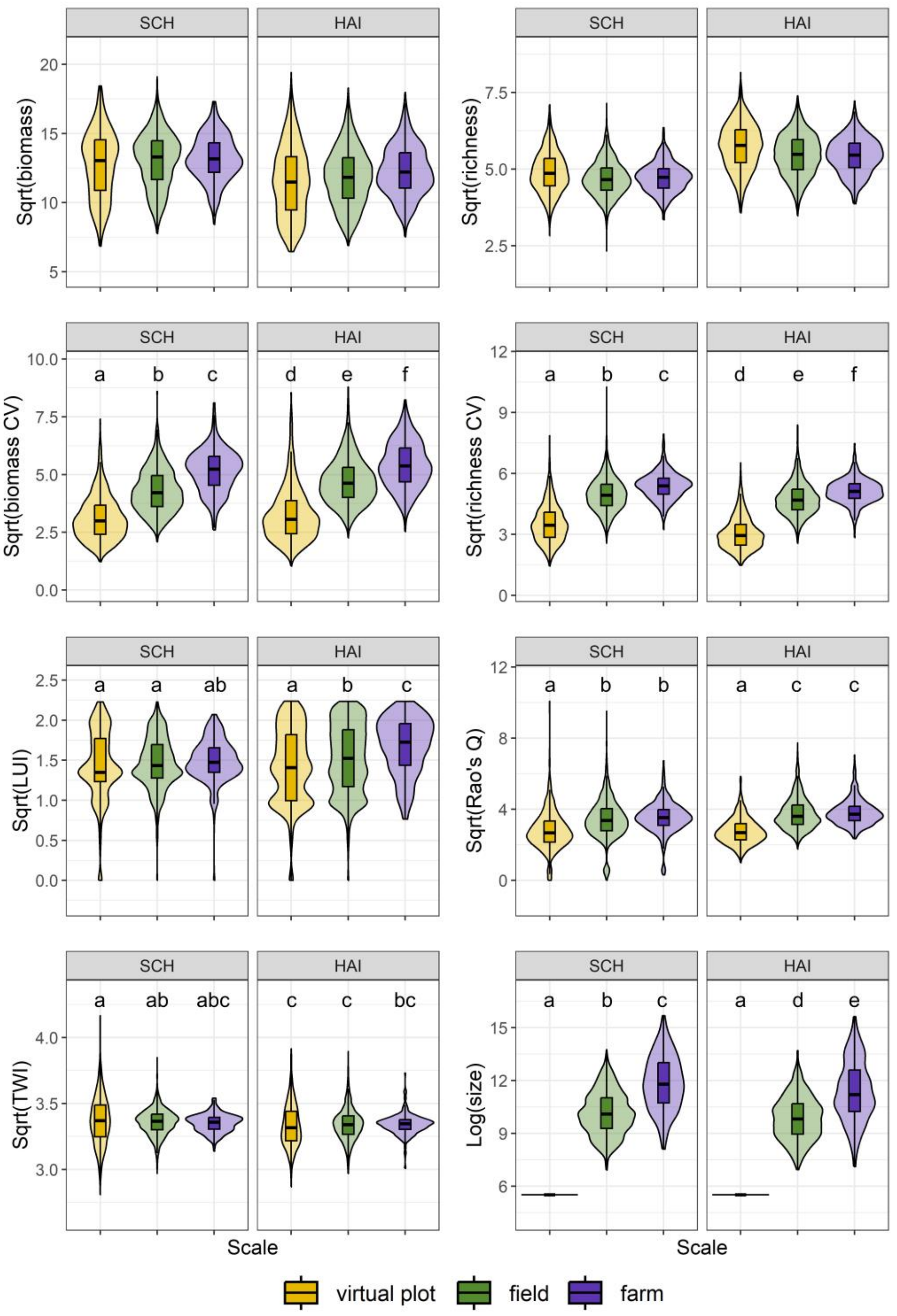


**Fig. S4** Differences between the spatial scale and region (SCH: Schorfheide-Chorin, HAI: Hainich-Dün) in response to biomass and its spatial variability (biomass CV), species richness and its spatial variability (richness CV), land-use intensity (LUI), spectral heterogeneity (Rao's Q), topographic wetness index (TWI) and size of the respective spatial unit. Different letters indicate significant differences between the interactions of spatial scale and exploratory region for the respective variable, which derived from post-hoc Tukey tests. There was no interaction detected on biomass ($F_{2,9731}$ = 2.34, p-value > 0.05) and species richness ($F_{2,9731}$ = 2.55, p-value > 0.05) and thus no post-hoc Tukey test was applied. The interaction between spatial scale and exploratory region was significant for biomass CV ($F_{2,9731}$ = 14.69, p-value < 0.001***), richness CV ($F_{2,9731}$ = 25.26, p-value < 0.001***), LUI ($F_{2,9731}$ = 18.60, p-value < 0.001***), Rao's Q ($F_{2,9731}$ = 28.68, p-value < 0.001***), TWI ($F_{2,9731}$ = 6.62, p-value < 0.01**) and size ($F_{2,9731}$ = 19.95, p-value < 0.001***)

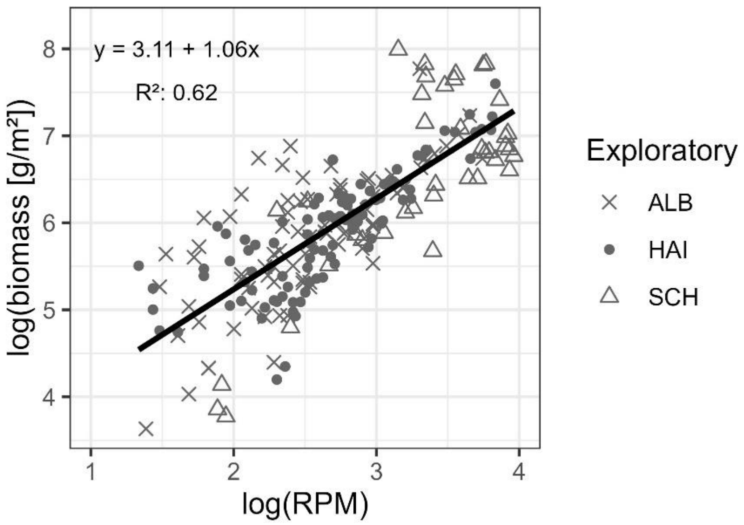


**Fig. S5** Calibration curve for Rising Plate Meter (RPM) measurements and biomass cuttings in grassland plots of the biodiversity exploratories. Field data from all three regions of the biodiversity exploratory are included (ALB: Schwäbische Alb, HAI: Hainich-Dün, SCH: Schorfheide-Chorin)

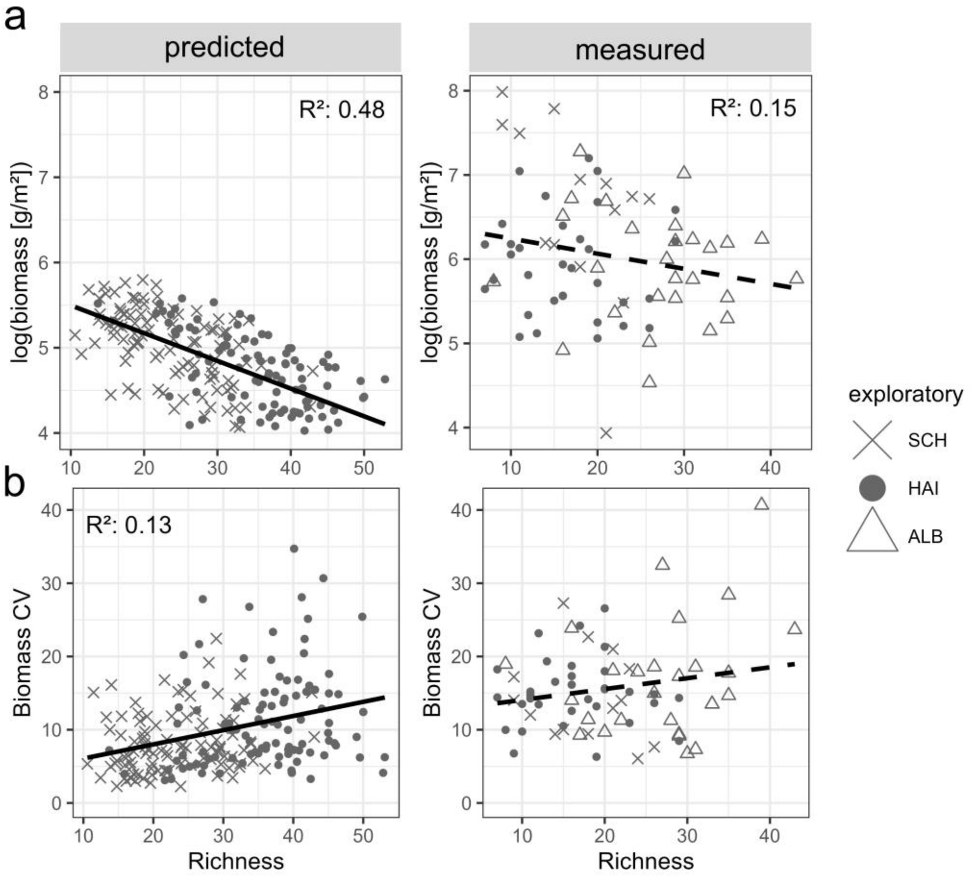


**Fig. S6** Relationship between (a) species richness and biomass as well as between (b) species richness and the spatial variability of biomass (biomass CV) based on predicted and measured data in grassland plots of the biodiversity exploratories. Significant relationships are indicated with solid lines, while non-significant relationships are represented as dashed lines. Different shapes of points indicate the different exploratory regions Schorfheide-Chorin (SCH), Hainich-Dün (HAI) and Schwäbische Alb (ALB). Although data from multiple years are included in the model, individual years are not explicitly shown. Note that predicted data represents species richness sampled in 16 m² and biomass CV was calculated based on about 25 pixels, whereas species richness of measured data represents 3 m² and biomass CV refers to 10 observations per plot. The multiple linear regression using measured data with biomass CV as the response variable, was not significant, therefore no $R^2$ is reported (see Table S18)

### Supplementary tables

**Table S1** Count of polygons at virtual plot, field and farm scale for each of the four subsets based on region (SCH: Schorfheide-Chorin, HAI: Hainich-Dün) and year (2020, 2021) after the filtering process and the removal of missing observations

| | | SCH 2020 | SCH 2021 | HAI 2020 | HAI 2021 |
|---|---|---|---|---|---|
| # Virtual plots | | 516 | 515 | 636 | 628 |
| Fields per grassland type | Meadows | 389 | 371 | 150 | 165 |
| | Pastures | 45 | 40 | 358 | 328 |
| | Mown pastures | 785 | 790 | 1461 | 1468 |
| | Other grasslands | 85 | 82 | - | - |
| # Fields | | 1304 | 1283 | 1969 | 1961 |
| # Farms | | 184 | 190 | 277 | 274 |

**Table S2** Mean and standard deviation of all aggregated pixel values for each respective variable and spatial scale analysed in this study. Biomass values are expressed in g $m^{-2}$ and the size of virtual plots, fields and farms are expressed in ha

| | Variable | SCH 2020 | SCH 2021 | HAI 2020 | HAI 2021 |
|---|---|---|---|---|---|
| **Virtual plot scale** | Biomass | 152.28 ± 57.39 | 185.39 ± 60.84 | 132.36 ± 63.06 | 146.11 ± 57.29 |
| | Biomass CV | 11.48 ± 7.24 | 9.9 ± 6.53 | 13.47 ± 10.62 | 10.7 ± 7.75 |
| | Richness | 24.7 ± 6.31 | 24.33 ± 6.95 | 32.19 ± 8.3 | 34.69 ± 8.79 |
| | Richness CV | 12.19 ± 6.16 | 14.52 ± 7.87 | 9.39 ± 5.13 | 10.55 ± 5.82 |
| | LUI | 2.29 ± 1.19 | 2.24 ± 1.16 | 2.28 ± 1.32 | 2.25 ± 1.27 |
| | Spectral heterogeneity | 10.01 ± 6.43 | 7.84 ± 8.25 | 8.83 ± 4.3 | 7.58 ± 4.59 |
| | TWI | 11.4 ± 1.25 | 11.4 ± 1.26 | 11.17 ± 1.08 | 11.11 ± 1.05 |
| | Size | 0.025 ± 0 | 0.025 ± 0 | 0.025 ± 0 | 0.025 ± 0 |
| **Field scale** | Biomass | 161.11 ± 47.28 | 188.44 ± 49.96 | 138.91 ± 53.07 | 150.11 ± 44.77 |
| | Biomass CV | 20.43 ± 8.73 | 18.29 ± 8.13 | 25.44 ± 10.71 | 20.84 ± 8.08 |
| | Richness | 22.11 ± 5.03 | 22.55 ± 5.56 | 29.22 ± 7.21 | 31.52 ± 7.26 |
| | Richness CV | 23.97 ± 7.30 | 26.45 ± 8.90 | 22.33 ± 7.64 | 24.09 ± 8.29 |
| | LUI | 2.25 ± 0.92 | 2.24 ± 0.91 | 2.49 ± 1.28 | 2.50 ± 1.28 |
| | Spectral heterogeneity | 14.04 ± 7.07 | 11.77 ± 8.50 | 15.73 ± 7.14 | 13.97 ± 7.29 |
| | TWI | 11.29 ± 0.64 | 11.29 ± 0.65 | 11.17 ± 0.78 | 11.18 ± 0.78 |
| | Size | 5.35 ± 8.16 | 5.44 ± 8.27 | 4.08 ± 6.82 | 4.05 ± 6.79 |
| **Farm scale** | Biomass | 162.78 ± 40.38 | 190.9 ± 39.56 | 149.95 ± 47.93 | 158.99 ± 41.06 |
| | Biomass CV | 28.81 ± 8.84 | 26.17 ± 9.84 | 34.08 ± 12.29 | 26.52 ± 8.1 |
| | Richness | 22.55 ± 4.11 | 22.52 ± 4.83 | 28.42 ± 5.88 | 31.44 ± 6.29 |
| | Richness CV | 27.66 ± 5.56 | 31.17 ± 7.97 | 25.76 ± 6.02 | 27.53 ± 6.19 |
| | LUI | 2.29 ± 0.72 | 2.31 ± 0.77 | 2.92 ± 1.11 | 2.95 ± 1.12 |
| | Spectral heterogeneity | 14.56 ± 5.27 | 11.98 ± 6.46 | 15.83 ± 5.92 | 14.25 ± 5.95 |
| | TWI | 11.25 ± 0.45 | 11.23 ± 0.46 | 11.17 ± 0.51 | 11.18 ± 0.51 |
| | Size | 43.33 ± 80.65 | 41.89 ± 79.42 | 34.45 ± 75.17 | 34.72 ± 74.73 |

**Table S3** Hypothesised causal relationships for each path included in the a-priori conceptual structural equation model (SEM) testing direct and indirect land-use intensity (LUI) effects on the spatial variability of biomass (biomass CV). Bold marked paths refer to the first and second hypothesis of this study

| **Pathway** | **Hypothesised ecological relationships based on literature** |
|---|---|
| ***Richness* (α) ~ *LUI*** | High LUI negatively impacts plant species richness (Wesche et al. 2012; Beckmann et al. 2019). Particularly, specialist species with narrow niche widths are threatened due to increased competition (Busch et al. 2019), caused by intensification. However, along the entire gradient of productivity a unimodal distribution of diversity is reported (Fraser et al. 2015). |
| Richness (α) ~ Spectral heterogeneity | Spectral variation is caused by spatial habitat heterogeneity (spectral variation hypothesis of Palmer et al. (2002) and was reported to be both positively (Palmer et al. 2002; Wang et al. 2018) or negatively correlated with species richness (Rossi et al. 2022). An overall positive relationship is assumed, based on the assumption that habitat heterogeneity provides niches and is thus a main driver of increasing species richness (Stein et al. 2014; Maskell et al. 2019). |
| Richness (α) ~ Topographic wetness index | Soil moisture is influenced by topography and plays an important role at local and regional scales for species richness in dry or wet grasslands and was reported to be negatively correlated (Moeslund et al. 2013). |
| ***Richness CV* (α) ~ *LUI*** | Intensive land use leads to a spatial biotic homogenisation (decreased beta-diversity) of plant communities (Gossner et al. 2016; Liu et al. 2018). The variability of alpha diversity through space is thus also assumed to be reduced. |
| Richness CV (α) ~ Spectral heterogeneity | Spectral heterogeneity as a proxy for environmental heterogeneity (Rocchini et al. 2010) does not only has a positive effect on average species richness within a spatial extent, but also on the variability of alpha diversity through space since the influence of species turnover increases with spatial scale (Stein et al. 2014). |
| Richness CV (α) ~ Size | With increasing field size, the standard deviation becomes higher, leading to an increase of richness CV since the standard deviation is the numerator in the equation for the coefficient of variation (CV) (Taylor et al. 1999). |
| Richness CV (α) ~ Richness (α) | A greater species pool within a field leads to lower spatial variability of species richness, especially with the occurrence of more competitive species (Ives and Hughes 2002). Moreover, as reported for the relationship of biomass CV and biomass, there is an immanent negative correlation between the mean and coefficient of variation (CV) of the same variable since the mean is the denominator in the equation for the CV (Taylor et al. 1999). |
| ***Biomass CV* ~ *Richness* (α)** | Plant species richness is expected to stabilise the spatial variability of biomass (Daleo et al. 2023), which is explained by diversity acting as biological insurance through space that enables to provide ecosystem functions even when some species go extinct (Loreau et al. 2021). |

| | |
|---|---|
| ***Biomass CV ~ Richness CV* (α)** | There is a negative relationship between species covariation and the spatial variability of biomass production (Daleo et al. 2023) as well as a positive effect between the spatial synchrony of species richness in relation to the temporal stability of biomass (Walter et al. 2021). Similarly, it is assumed that the higher the spatial variability of species richness, the lower the spatial variability of biomass due to enhanced spatial insurance effects that protect against environmental fluctuations (Loreau et al. 2021). |
| Biomass CV ~ LUI | LUI decreases the stability of ecosystem's productivity (Canelas and Pereira 2022) and vice versa, increases spatial biomass variability. |
| Biomass CV ~ Spectral heterogeneity | Spatial variability of biomass increases with higher environmental heterogeneity (Daleo et al. 2023), thereby also with increasing spectral heterogeneity as a proxy for environmental heterogeneity (Rocchini et al. 2010). |
| Biomass CV ~ Size | With increasing field size, the standard deviation of aggregated pixels becomes higher, leading to an increase of biomass CV since the standard deviation is the numerator in the equation for the coefficient of variation (CV) (Taylor et al. 1999). |
| Biomass CV ~ Biomass | Variability in biomass production and average biomass is negatively correlated since the mean is included in the denominator of the equation for the coefficient of variation (CV) (Taylor et al. 1999). |
| Biomass ~ LUI | Conventional land-use intensification is increasing biomass productivity, e.g. through the application of fertilisation, to maximise the yield within grassland fields (Beckmann et al. 2019). |
| Biomass ~ Spectral heterogeneity | Based on the positive spectral variation hypothesis (Palmer et al. 2002), the opposite of a negative correlation between spectral heterogeneity and biomass is assumed since highly productive grasslands are usually low in species diversity (Andraczek et al. 2023) and are therefore assumed to be more homogeneous. |
| Biomass ~ Topographic wetness index | Productive plant species prefer more nitrogen rich soils, which are usually associated with more wetter areas (Moeslund et al. 2013). Therefore, biomass is expected to be higher with increasing TWI. |
| Biomass ~ Richness (α) | In productive landscapes, the relationship between biomass and plant species richness was found to be negative, but with decreasing LUI the relationship can even change towards positive (Andraczek et al. 2023). |

**Table S4** Model fit statistics and standardised path coefficients for each of the four piecewise structural equation models (pSEM) testing direct and indirect land-use intensity (LUI) effects on the spatial variability of biomass (biomass CV). Bold standardised estimates are related to the first and second hypothesis. The four pSEMs are based on different subsets regarding region (SCH: Schorfheide-Chorin, HAI: Hainich-Dün) and year. Species richness (richness) and its spatial variability (richness CV) represent mediators. Standard estimates of each path are shown with according significance levels (p<0.05*, p<0.01**, p<0.001***). Note that standardised estimates cannot be directly compared in terms of units since variables were transformed differently (see Table S5-S8), models tested different data subsets and the structure of pSEMs was slightly different

| **Model fit statistics** | | **SCH 2020** | **SCH 2021** | **HAI 2020** | **HAI 2021** |
|---|---|---|---|---|---|
| Fisher's C | | 9.190 | 10.559 | 10.526 | 6.502 |
| p-value | | 0.163 | 0.103 | 0.104 | 0.165 |
| Degrees of freedom | | 6 | 6 | 6 | 4 |
| AIC (d-sep) | | 11,665.72 | 19,257.43 | 9,361.316 | 8,627.08 |
| Sample size | | 1304 | 1283 | 1969 | 1961 |
| **Response** | **Predictor** | | | | |
| Richness | **LUI** | **-0.20*** ** | **-0.15*** ** | **-0.29*** ** | **-0.18*** ** |
| | Spectral heterogeneity | 0.07* | -0.08** | -0.14*** | -0.17*** |
| | TWI | -0.03 | -0.08** | -0.20*** | -0.18*** |
| | Size | 0.24*** | 0.19*** | 0.16*** | 0.13*** |
| Biomass | LUI | 0.09*** | 0.03 | 0.11*** | 0.13*** |
| | Richness | -0.59*** | -0.40*** | -0.66*** | -0.55*** |
| | Spectral heterogeneity | -0.20*** | -0.15*** | -0.05** | -0.05** |
| | TWI | 0.02 | -0.02 | 0.09*** | 0.12*** |
| Richness CV | **LUI** | **-0.08** ** | **-0.13*** ** | **-0.06** ** | **-0.03** |
| | Richness | -0.21*** | -0.31*** | -0.52*** | -0.53*** |
| | Spectral heterogeneity | 0.49*** | 0.27*** | 0.15*** | 0.14*** |
| | Size | 0.17*** | 0.14*** | -0.07** | -0.11*** |
| | TWI | - | - | -0.04* | -0.06** |
| | Biomass | - | - | - | -0.05* |
| Biomass CV | **Richness** | **0.11*** ** | **0.07** ** | **0.44*** ** | **0.02** |
| | **Richness CV** | **0.20*** ** | **0.28*** ** | **0.29*** ** | **0.21*** ** |
| | LUI | 0.02 | 0.02 | 0.03 | -0.03 |
| | Spectral heterogeneity | 0.31*** | 0.15*** | 0.16*** | 0.25*** |
| | Biomass | -0.32*** | -0.47*** | -0.15*** | -0.46*** |
| | Size | 0.36*** | 0.24*** | 0.29*** | 0.29*** |
| | TWI | -0.06** | -0.04 | - | - |

**Table S5** Model fit statistics of piecewise structural equation model (pSEM) testing direct and indirect land-use intensity (LUI) effects on the spatial variability of biomass (biomass CV) in Schorfheide-Chorin for the year 2020 at the field scale. Adjusted $R^2$ is shown for each response variable which was included in the model

| Response | Predictor | Estimate | Std. Error | DF | p-value | Std. Estimate |
|---|---|---|---|---|---|---|
| sqrt(richness) $R^2$ = 0.10 | sqrt(LUI) | -0.34 | 0.05 | 1299 | <0.001*** | -0.20 |
| | log(Rao's Q) | 0.08 | 0.03 | 1299 | <0.05* | 0.07 |
| | sqrt(TWI) | -0.19 | 0.15 | 1299 | 0.20 | -0.03 |
| | log(size) | 0.10 | 0.01 | 1299 | <0.001*** | 0.24 |
| sqrt(biomass) $R^2$ = 0.44 | sqrt(LUI) | 0.52 | 0.13 | 1299 | <0.001*** | 0.09 |
| | sqrt(richness) | -2.12 | 0.08 | 1299 | <0.001*** | -0.59 |
| | log(Rao's Q) | -0.8 | 0.08 | 1299 | <0.001*** | -0.20 |
| | sqrt(TWI) | 0.34 | 0.41 | 1299 | 0.42 | 0.02 |
| sqrt(richness CV) $R^2$ = 0.26 | sqrt(richness) | -0.3 | 0.04 | 1299 | <0.001*** | -0.21 |
| | log(Rao's Q) | 0.76 | 0.04 | 1299 | <0.001*** | 0.49 |
| | sqrt(LUI) | -0.18 | 0.06 | 1299 | <0.01** | -0.08 |
| | log(size) | 0.10 | 0.02 | 1299 | <0.001*** | 0.17 |
| sqrt(biomass CV) $R^2$ = 0.52 | log(Rao's Q) | 0.61 | 0.05 | 1296 | <0.001*** | 0.31 |
| | sqrt(richness) | 0.21 | 0.05 | 1296 | <0.001*** | 0.11 |
| | sqrt(LUI) | 0.05 | 0.06 | 1296 | 0.44 | 0.02 |
| | sqrt(richness) | 0.25 | 0.03 | 1296 | <0.001*** | 0.20 |
| | sqrt(biomass) | -0.16 | 0.01 | 1296 | <0.001*** | -0.32 |
| | log(size) | 0.28 | 0.02 | 1296 | <0.001*** | 0.36 |
| | sqrt(TWI) | -0.63 | 0.19 | 1296 | <0.01** | -0.06 |

**Table S6** Model fit statistics of piecewise structural equation model (pSEM) testing direct and indirect land-use intensity (LUI) effects on the spatial variability of biomass (biomass CV) in Schorfheide-Chorin for the year 2021 at the field scale. Adjusted $R^2$ is shown for each response variable which was included in the model

| Response | Predictor | Estimate | Std. Error | DF | p-value | Std. Estimate |
|---|---|---|---|---|---|---|
| sqrt(richness) $R^2$ = 0.07 | sqrt(LUI) | -0.27 | 0.05 | 1278 | <0.001*** | -0.15 |
| | sqrt(Rao's Q) | -0.04 | 0.01 | 1278 | <0.01** | -0.08 |
| | sqrt(TWI) | -0.47 | 0.16 | 1278 | <0.01** | -0.08 |
| | log(size) | 0.09 | 0.01 | 1278 | <0.001*** | 0.19 |
| Biomass $R^2$ = 0.18 | sqrt(LUI) | 4.57 | 4.09 | 1278 | 0.26 | 0.03 |
| | sqrt(richness) | -34.5 | 2.23 | 1278 | <0.001*** | -0.40 |
| | sqrt(Rao's Q) | -6.23 | 1.05 | 1278 | <0.001*** | -0.15 |
| | sqrt(TWI) | -9.72 | 13.25 | 1278 | 0.46 | -0.02 |
| sqrt(richness CV) $R^2$ = 0.18 | sqrt(richness) | -0.45 | 0.04 | 1278 | <0.001*** | -0.31 |
| | sqrt(Rao's Q) | 0.18 | 0.02 | 1278 | <0.001*** | 0.27 |
| | sqrt(LUI) | -0.34 | 0.07 | 1278 | <0.001*** | -0.13 |
| | log(size) | 0.09 | 0.02 | 1278 | <0.001*** | 0.14 |
| log(biomass CV) $R^2$ = 0.45 | sqrt(Rao's Q) | 0.05 | 0.01 | 1275 | <0.001*** | 0.15 |
| | sqrt(richness) | 0.05 | 0.02 | 1275 | <0.01** | 0.07 |
| | sqrt(LUI) | 0.02 | 0.03 | 1275 | 0.43 | 0.02 |
| | sqrt(richness CV) | 0.14 | 0.01 | 1275 | <0.001*** | 0.28 |
| | biomass | 0.00 | 0.00 | 1275 | <0.001*** | -0.47 |
| | log(size) | 0.08 | 0.01 | 1275 | <0.001*** | 0.24 |
| | sqrt(TWI) | -0.16 | 0.09 | 1275 | 0.09 | -0.04 |

**Table S7** Model fit statistics of piecewise structural equation model (pSEM) testing direct and indirect land-use intensity (LUI) effects on the spatial variability of biomass (biomass CV) in Hainich-Dün for the year 2020 at the field scale. Adjusted $R^2$ is shown for each response variable which was included in the model

| Response | Predictor | Estimate | Std. Error | DF | p-value | Std. Estimate |
|---|---|---|---|---|---|---|
| sqrt(richness) $R^2$ = 0.19 | sqrt(LUI) | -0.46 | 0.03 | 1964 | <0.001*** | -0.29 |
| | sqrt(Rao's Q) | -0.12 | 0.02 | 1964 | <0.001*** | -0.14 |
| | sqrt(TWI) | -1.15 | 0.12 | 1964 | <0.001*** | -0.20 |
| | log(size) | 0.09 | 0.01 | 1964 | <0.001*** | 0.16 |
| log(biomass) $R^2$ = 0.51 | sqrt(LUI) | 0.10 | 0.02 | 1964 | <0.001*** | 0.11 |
| | sqrt(richness) | -0.38 | 0.01 | 1964 | <0.001*** | -0.66 |
| | sqrt(Rao's Q) | -0.02 | 0.01 | 1964 | <0.01** | -0.05 |
| | sqrt(TWI) | 0.28 | 0.05 | 1964 | <0.001*** | 0.09 |
| sqrt(richness CV) $R^2$ = 0.34 | sqrt(richness) | -0.6 | 0.02 | 1963 | <0.001*** | -0.52 |
| | sqrt(Rao's Q) | 0.14 | 0.02 | 1963 | <0.001*** | 0.15 |
| | sqrt(LUI) | -0.11 | 0.04 | 1963 | <0.01** | -0.06 |
| | log(size) | -0.04 | 0.01 | 1963 | <0.01** | -0.07 |
| | sqrt(TWI) | -0.28 | 0.13 | 1963 | <0.05* | -0.04 |
| log(biomass CV) $R^2$ = 0.29 | sqrt(Rao's Q) | 0.08 | 0.01 | 1962 | <0.001*** | 0.16 |
| | sqrt(richness) | 0.27 | 0.02 | 1962 | <0.001*** | 0.44 |
| | sqrt(LUI) | 0.03 | 0.02 | 1962 | 0.13 | 0.03 |
| | sqrt(richness CV) | 0.16 | 0.01 | 1962 | <0.001*** | 0.29 |
| | log(biomass) | -0.16 | 0.03 | 1962 | <0.001*** | -0.15 |
| | log(size) | 0.10 | 0.01 | 1962 | <0.001*** | 0.29 |

**Table S8** Model fit statistics of piecewise structural equation model (pSEM) testing direct and indirect land-use intensity (LUI) effects on the spatial variability of biomass (biomass CV) in Hainich-Dün for the year 2021 at the field scale. Adjusted $R^2$ is shown for each response variable which was included in the model

| Response | Predictor | Estimate | Std. Error | DF | p-value | Std. Estimate |
|---|---|---|---|---|---|---|
| sqrt(richness) $R^2$ = 0.13 | sqrt(LUI) | -0.28 | 0.03 | 1956 | <0.001*** | -0.18 |
| | sqrt(Rao's Q) | -0.12 | 0.02 | 1956 | <0.001*** | -0.17 |
| | sqrt(TWI) | -0.99 | 0.12 | 1956 | <0.001*** | -0.18 |
| | log(size) | 0.07 | 0.01 | 1956 | <0.001*** | 0.13 |
| log(biomass) $R^2$ = 0.38 | sqrt(LUI) | 0.09 | 0.01 | 1956 | <0.001*** | 0.13 |
| | sqrt(richness) | -0.26 | 0.01 | 1956 | <0.001*** | -0.55 |
| | sqrt(Rao's Q) | -0.02 | 0.01 | 1956 | <0.01** | -0.05 |
| | sqrt(TWI) | 0.31 | 0.05 | 1956 | <0.001*** | 0.12 |
| sqrt(richness CV) $R^2$ = 0.34 | sqrt(richness) | -0.66 | 0.03 | 1954 | <0.001*** | -0.53 |
| | sqrt(Rao's Q) | 0.12 | 0.02 | 1954 | <0.001*** | 0.14 |
| | sqrt(LUI) | -0.05 | 0.04 | 1954 | 0.18 | -0.03 |
| | log(size) | -0.07 | 0.01 | 1954 | <0.001*** | -0.11 |
| | sqrt(TWI) | -0.41 | 0.13 | 1954 | <0.01** | -0.06 |
| | log(biomass) | -0.14 | 0.06 | 1954 | <0.05* | -0.05 |
| log(biomass CV) $R^2$ = 0.31 | sqrt(Rao's Q) | 0.11 | 0.01 | 1954 | <0.001*** | 0.25 |
| | sqrt(richness) | 0.01 | 0.02 | 1954 | 0.52 | 0.02 |
| | sqrt(LUI) | -0.03 | 0.02 | 1954 | 0.10 | -0.03 |
| | sqrt(richness CV) | 0.10 | 0.01 | 1954 | <0.001*** | 0.21 |
| | log(biomass) | -0.57 | 0.03 | 1954 | <0.001*** | -0.46 |
| | log(size) | 0.09 | 0.01 | 1954 | <0.001*** | 0.29 |

**Table S9** Model fit statistics of the multiple linear regression testing the relationship between species richness and biomass across spatial scales and different land-use intensities (LUI) in the region Schorfheide-Chorin for the year 2020. All continuous variables were square root transformed, except the variable size. A three-way interaction between species richness, spatial scale and LUI category (low, medium, high) was tested. Interaction terms are indicated with a colon. Adjusted $R^2$ = 0.46, $F_{20,1983}$ = 87.43, p-value <0.001***

| Predictor | Estimate | SE | 95% CI | t-value | p-value |
|---|---|---|---|---|---|
| (Intercept) | 23.89 | 1.21 | [21.52, 26.27] | 19.73 | <0.001*** |
| sqrt(mean_rich) | -2.3 | 0.15 | [-2.6, -2.01] | -15.44 | <0.001*** |
| scalefield | -1.07 | 0.94 | [-2.91, 0.78] | -1.13 | 0.257 |
| scalefarm | 3.43 | 2.33 | [-1.15, 8.01] | 1.47 | 0.142 |
| LUImedium | -0.38 | 1.2 | [-2.73, 1.97] | -0.32 | 0.752 |
| LUIhigh | -1.79 | 1.33 | [-4.41, 0.82] | -1.35 | 0.178 |
| sqrt(mean_RaoQ) | -0.49 | 0.04 | [-0.56, -0.41] | -12.28 | <0.001*** |
| sqrt(mean_TWI) | 0.3 | 0.27 | [-0.23, 0.83] | 1.11 | 0.267 |
| log(size) | -0.03 | 0.03 | [-0.09, 0.04] | -0.79 | 0.43 |
| sqrt(mean_rich):scalefield | 0.23 | 0.19 | [-0.14, 0.6] | 1.2 | 0.23 |
| sqrt(mean_rich):scalefarm | -0.59 | 0.48 | [-1.52, 0.35] | -1.23 | 0.22 |
| sqrt(mean_rich):LUImedium | 0.03 | 0.24 | [-0.44, 0.5] | 0.11 | 0.909 |
| sqrt(mean_rich):LUIhigh | 0.38 | 0.27 | [-0.16, 0.91] | 1.39 | 0.166 |
| scalefield:LUImedium | 1.57 | 1.44 | [-1.25, 4.38] | 1.09 | 0.276 |
| scalefarm:LUImedium | 0.36 | 2.95 | [-5.43, 6.15] | 0.12 | 0.903 |
| scalefield:LUIhigh | 0.87 | 1.79 | [-2.65, 4.39] | 0.49 | 0.627 |
| scalefarm:LUIhigh | -13.64 | 6.18 | [-25.76, -1.51] | -2.21 | <0.05* |
| sqrt(mean_rich):scalefield:LUI medium | -0.21 | 0.29 | [-0.79, 0.36] | -0.73 | 0.468 |
| sqrt(mean_rich):scalefarm:LUI medium | -0.04 | 0.61 | [-1.24, 1.15] | -0.07 | 0.943 |
| sqrt(mean_rich):scalefield:LUI high | -0.09 | 0.38 | [-0.83, 0.65] | -0.23 | 0.815 |
| sqrt(mean_rich):scalefarm:LUI high | 2.91 | 1.34 | [0.28, 5.53] | 2.17 | <0.05* |

**Table S10** Model fit statistics of the multiple linear regression testing the relationship between species richness and biomass across spatial scales and different land-use intensities (LUI) in the region Schorfheide-Chorin for the year 2021. All continuous variables were square root transformed, except the variable size. A three-way interaction between species richness, spatial scale and LUI category (low, medium, high) was tested. Interaction terms are indicated with a colon. Adjusted $R^2$ = 0.19, $F_{20,1967}$ = 23.66, p-value <0.001***

| Predictor | Estimate | SE | 95% CI | t-value | p-value |
|---|---|---|---|---|---|
| (Intercept) | 21.64 | 1.41 | [18.88, 24.4] | 15.38 | <0.001*** |
| sqrt(mean_rich) | -1.27 | 0.16 | [-1.58, -0.96] | -8.09 | <0.001*** |
| scalefield | 0.17 | 1.03 | [-1.84, 2.18] | 0.17 | 0.867 |
| scalefarm | 1.11 | 2.49 | [-3.76, 5.99] | 0.45 | 0.655 |
| LUImedium | 0.88 | 1.41 | [-1.89, 3.65] | 0.62 | 0.535 |
| LUIhigh | 1.12 | 1.39 | [-1.61, 3.86] | 0.81 | 0.42 |
| sqrt(mean_RaoQ) | -0.23 | 0.03 | [-0.3, -0.16] | -6.75 | <0.001*** |
| sqrt(mean_TWI) | -0.45 | 0.33 | [-1.09, 0.19] | -1.38 | 0.168 |
| log(size) | -0.04 | 0.04 | [-0.11, 0.04] | -0.96 | 0.338 |
| sqrt(mean_rich):scalefield | -0.02 | 0.21 | [-0.42, 0.39] | -0.08 | 0.938 |
| sqrt(mean_rich):scalefarm | -0.13 | 0.51 | [-1.14, 0.87] | -0.26 | 0.794 |
| sqrt(mean_rich):LUImedium | -0.19 | 0.29 | [-0.75, 0.38] | -0.66 | 0.512 |
| sqrt(mean_rich):LUIhigh | -0.27 | 0.29 | [-0.83, 0.3] | -0.93 | 0.355 |
| scalefield:LUImedium | 0.53 | 1.67 | [-2.75, 3.82] | 0.32 | 0.749 |
| scalefarm:LUImedium | -0.06 | 3.2 | [-6.34, 6.21] | -0.02 | 0.985 |
| scalefield:LUIhigh | -4.55 | 1.84 | [-8.16, -0.94] | -2.47 | <0.05* |
| scalefarm:LUIhigh | 0.67 | 4.1 | [-7.37, 8.72] | 0.16 | 0.869 |
| sqrt(mean_rich):scalefield:LUI medium | -0.05 | 0.34 | [-0.72, 0.62] | -0.15 | 0.884 |
| sqrt(mean_rich):scalefarm:LUI medium | 0.03 | 0.66 | [-1.27, 1.32] | 0.04 | 0.967 |
| sqrt(mean_rich):scalefield:LUI high | 1.04 | 0.39 | [0.28, 1.79] | 2.69 | <0.01** |
| sqrt(mean_rich):scalefarm:LUI high | -0.18 | 0.87 | [-1.89, 1.53] | -0.2 | 0.839 |

**Table S11** Model fit statistics of the multiple linear regression testing the relationship between species richness and biomass across spatial scales and different land-use intensities (LUI) in the region Hainich-Dün for the year 2020. All continuous variables were square root transformed, except the variable size. A three-way interaction between species richness, spatial scale and LUI category (low, medium, high) was tested. Interaction terms are indicated with a colon. Adjusted $R^2$ = 0.51, $F_{20,2861}$ = 148.7, p-value <0.001***

| Predictor | Estimate | SE | 95% CI | t-value | p-value |
|---|---|---|---|---|---|
| (Intercept) | 19.35 | 1.17 | [17.06, 21.64] | 16.57 | <0.001*** |
| sqrt(mean_rich) | -2.34 | 0.13 | [-2.6, -2.08] | -17.42 | <0.001*** |
| scalefield | -0.98 | 0.94 | [-2.82, 0.87] | -1.04 | 0.3 |
| scalefarm | 3.7 | 2.27 | [-0.75, 8.14] | 1.63 | 0.103 |
| LUImedium | 1.53 | 1.27 | [-0.97, 4.03] | 1.2 | 0.229 |
| LUIhigh | -0.6 | 1.2 | [-2.96, 1.76] | -0.5 | 0.619 |
| sqrt(mean_RaoQ) | -0.17 | 0.04 | [-0.25, -0.08] | -3.9 | <0.001*** |
| sqrt(mean_TWI) | 1.56 | 0.25 | [1.07, 2.05] | 6.25 | <0.001*** |
| log(size) | 0 | 0.03 | [-0.06, 0.06] | 0.02 | 0.987 |
| sqrt(mean_rich):scalefield | 0.17 | 0.16 | [-0.15, 0.49] | 1.04 | 0.3 |
| sqrt(mean_rich):scalefarm | -0.6 | 0.41 | [-1.41, 0.2] | -1.46 | 0.143 |
| sqrt(mean_rich):LUImedium | -0.2 | 0.22 | [-0.64, 0.23] | -0.92 | 0.358 |
| sqrt(mean_rich):LUIhigh | 0.22 | 0.22 | [-0.2, 0.64] | 1.03 | 0.303 |
| scalefield:LUImedium | -1.14 | 1.48 | [-4.04, 1.76] | -0.77 | 0.439 |
| scalefarm:LUImedium | -4.15 | 2.9 | [-9.83, 1.53] | -1.43 | 0.152 |
| scalefield:LUIhigh | 1.49 | 1.42 | [-1.29, 4.27] | 1.05 | 0.293 |
| scalefarm:LUIhigh | -0.41 | 2.88 | [-6.05, 5.23] | -0.14 | 0.887 |
| sqrt(mean_rich):scalefield:LUI medium | 0.2 | 0.26 | [-0.32, 0.71] | 0.75 | 0.453 |
| sqrt(mean_rich):scalefarm:LUI medium | 0.7 | 0.53 | [-0.34, 1.73] | 1.32 | 0.186 |
| sqrt(mean_rich):scalefield:LUI high | -0.28 | 0.26 | [-0.78, 0.22] | -1.09 | 0.276 |
| sqrt(mean_rich):scalefarm:LUI high | 0.02 | 0.53 | [-1.02, 1.05] | 0.03 | 0.975 |

**Table S12** Model fit statistics of the multiple linear regression testing the relationship between species richness and biomass across spatial scales and different land-use intensities (LUI) in the region Hainich-Dün for the year 2021. All continuous variables were square root transformed, except the variable size. A three-way interaction between species richness, spatial scale and LUI category (low, medium, high) was tested. Interaction terms are indicated with a colon. Adjusted $R^2$ = 0.39, $F_{20,2842}$ = 92.82, p-value <0.001***

| Predictor | Estimate | SE | 95% CI | t-value | p-value |
|---|---|---|---|---|---|
| (Intercept) | 16.74 | 1.11 | [14.57, 18.92] | 15.08 | <0.001*** |
| sqrt(mean_rich) | -1.85 | 0.13 | [-2.1, -1.6] | -14.44 | <0.001*** |
| scalefield | 0.07 | 0.94 | [-1.76, 1.91] | 0.08 | 0.938 |
| scalefarm | 3.19 | 2.1 | [-0.93, 7.31] | 1.52 | 0.129 |
| LUImedium | -2.32 | 1.12 | [-4.52, -0.13] | -2.08 | <0.05* |
| LUIhigh | -0.61 | 1.2 | [-2.97, 1.75] | -0.51 | 0.611 |
| sqrt(mean_RaoQ) | -0.21 | 0.04 | [-0.29, -0.14] | -5.76 | <0.001*** |
| sqrt(mean_TWI) | 1.8 | 0.23 | [1.34, 2.26] | 7.67 | <0.001*** |
| log(size) | -0.05 | 0.03 | [-0.1, 0.01] | -1.75 | 0.08 |
| sqrt(mean_rich):scalefield | 0.04 | 0.16 | [-0.27, 0.35] | 0.24 | 0.811 |
| sqrt(mean_rich):scalefarm | -0.42 | 0.36 | [-1.13, 0.29] | -1.15 | 0.25 |
| sqrt(mean_rich):LUImedium | 0.49 | 0.19 | [0.12, 0.86] | 2.62 | <0.01** |
| sqrt(mean_rich):LUIhigh | 0.2 | 0.21 | [-0.2, 0.61] | 0.98 | 0.329 |
| scalefield:LUImedium | 0.85 | 1.34 | [-1.77, 3.47] | 0.64 | 0.525 |
| scalefarm:LUImedium | 0.63 | 2.75 | [-4.77, 6.03] | 0.23 | 0.82 |
| scalefield:LUIhigh | -1.18 | 1.41 | [-3.93, 1.58] | -0.84 | 0.402 |
| scalefarm:LUIhigh | 0.34 | 2.7 | [-4.95, 5.63] | 0.13 | 0.9 |
| sqrt(mean_rich):scalefield:LUI medium | -0.19 | 0.23 | [-0.64, 0.26] | -0.82 | 0.413 |
| sqrt(mean_rich):scalefarm:LUI medium | -0.16 | 0.48 | [-1.1, 0.77] | -0.34 | 0.732 |
| sqrt(mean_rich):scalefield:LUI high | 0.2 | 0.24 | [-0.28, 0.68] | 0.83 | 0.407 |
| sqrt(mean_rich):scalefarm:LUI high | -0.14 | 0.47 | [-1.06, 0.79] | -0.29 | 0.774 |

**Table S13** Model fit statistics of the multiple linear regression testing the relationship between species richness and the spatial variability of biomass (biomass CV) across spatial scales and different land-use intensities (LUI) in the region Schorfheide-Chorin for the year 2020. All continuous variables were square root transformed, except the variable size. A three-way interaction between species richness, spatial scale and LUI category (low, medium, high) was tested. Interaction terms are indicated with a colon. Adjusted $R^2$ = 0.58, $F_{20,1983}$ = 139.60, p-value <0.001***

| Predictor | Estimate | SE | 95% CI | t-value | p-value |
|---|---|---|---|---|---|
| (Intercept) | 4.04 | 0.6 | [2.86, 5.22] | 6.71 | <0.001*** |
| sqrt(mean_rich) | 0.05 | 0.07 | [-0.09, 0.2] | 0.74 | 0.461 |
| scalefield | -2.54 | 0.47 | [-3.46, -1.62] | -5.43 | <0.001*** |
| scalefarm | -4.54 | 1.16 | [-6.81, -2.26] | -3.91 | <0.001*** |
| LUImedium | -1.25 | 0.6 | [-2.42, -0.08] | -2.1 | <0.05* |
| LUIhigh | 0.11 | 0.66 | [-1.19, 1.41] | 0.17 | 0.867 |
| sqrt(mean_RaoQ) | 0.52 | 0.02 | [0.48, 0.56] | 26.47 | <0.001*** |
| sqrt(mean_TWI) | -0.46 | 0.13 | [-0.73, -0.2] | -3.44 | <0.01** |
| log(size) | 0.3 | 0.02 | [0.27, 0.33] | 18.66 | <0.001*** |
| sqrt(mean_rich):scalefield | 0.45 | 0.09 | [0.27, 0.64] | 4.8 | <0.001*** |
| sqrt(mean_rich):scalefarm | 0.92 | 0.24 | [0.46, 1.39] | 3.88 | <0.001*** |
| sqrt(mean_rich):LUImedium | 0.26 | 0.12 | [0.02, 0.49] | 2.15 | <0.05* |
| sqrt(mean_rich):LUIhigh | 0 | 0.14 | [-0.27, 0.27] | 0 | 0.999 |
| scalefield:LUImedium | 1.35 | 0.71 | [-0.05, 2.75] | 1.89 | 0.059 |
| scalefarm:LUImedium | 3.04 | 1.47 | [0.16, 5.91] | 2.07 | <0.05* |
| scalefield:LUIhigh | 0.26 | 0.89 | [-1.49, 2.01] | 0.29 | 0.773 |
| scalefarm:LUIhigh | 3.89 | 3.07 | [-2.14, 9.91] | 1.27 | 0.206 |
| sqrt(mean_rich):scalefield:LUI medium | -0.3 | 0.15 | [-0.58, -0.01] | -2.05 | <0.05* |
| sqrt(mean_rich):scalefarm:LUI medium | -0.64 | 0.3 | [-1.23, -0.05] | -2.12 | <0.05* |
| sqrt(mean_rich):scalefield:LUI high | -0.11 | 0.19 | [-0.47, 0.26] | -0.56 | 0.573 |
| sqrt(mean_rich):scalefarm:LUI high | -0.9 | 0.67 | [-2.2, 0.41] | -1.35 | 0.178 |

**Table S14** Model fit statistics of the multiple linear regression testing the relationship between species richness and the spatial variability of biomass (biomass CV) across spatial scales and different land-use intensities (LUI) in the region Schorfheide-Chorin for the year 2021. All continuous variables were square root transformed, except the variable size. A three-way interaction between species richness, spatial scale and LUI category (low, medium, high) was tested. Interaction terms are indicated with a colon. Adjusted $R^2$ = 0.43, $F_{20,1967}$ = 76.99, p-value <0.001***

| Predictor | Estimate | SE | 95% CI | t-value | p-value |
|---|---|---|---|---|---|
| (Intercept) | 2.05 | 0.65 | [0.77, 3.33] | 3.13 | <0.01** |
| sqrt(mean_rich) | 0.25 | 0.07 | [0.11, 0.4] | 3.48 | <0.01** |
| scalefield | -0.27 | 0.48 | [-1.21, 0.67] | -0.57 | 0.571 |
| scalefarm | -0.2 | 1.16 | [-2.46, 2.07] | -0.17 | 0.865 |
| LUImedium | 0.47 | 0.66 | [-0.82, 1.76] | 0.72 | 0.474 |
| LUIhigh | 0.55 | 0.65 | [-0.72, 1.83] | 0.86 | 0.392 |
| sqrt(mean_RaoQ) | 0.24 | 0.02 | [0.21, 0.27] | 14.86 | <0.001*** |
| sqrt(mean_TWI) | -0.04 | 0.15 | [-0.33, 0.26] | -0.23 | 0.816 |
| log(size) | 0.2 | 0.02 | [0.17, 0.24] | 11.11 | <0.001*** |
| sqrt(mean_rich):scalefield | 0.09 | 0.1 | [-0.09, 0.28] | 0.98 | 0.325 |
| sqrt(mean_rich):scalefarm | 0.18 | 0.24 | [-0.29, 0.65] | 0.75 | 0.453 |
| sqrt(mean_rich):LUImedium | -0.1 | 0.13 | [-0.36, 0.16] | -0.74 | 0.461 |
| sqrt(mean_rich):LUIhigh | -0.14 | 0.13 | [-0.4, 0.12] | -1.04 | 0.298 |
| scalefield:LUImedium | -0.31 | 0.78 | [-1.83, 1.22] | -0.39 | 0.694 |
| scalefarm:LUImedium | -1.38 | 1.49 | [-4.3, 1.54] | -0.93 | 0.353 |
| scalefield:LUIhigh | 0.5 | 0.86 | [-1.18, 2.18] | 0.58 | 0.563 |
| scalefarm:LUIhigh | -1.13 | 1.91 | [-4.88, 2.61] | -0.59 | 0.552 |
| sqrt(mean_rich):scalefield:LUI medium | 0.03 | 0.16 | [-0.28, 0.35] | 0.22 | 0.828 |
| sqrt(mean_rich):scalefarm:LUI medium | 0.27 | 0.31 | [-0.33, 0.87] | 0.88 | 0.382 |
| sqrt(mean_rich):scalefield:LUI high | -0.1 | 0.18 | [-0.46, 0.25] | -0.58 | 0.56 |
| sqrt(mean_rich):scalefarm:LUI high | 0.27 | 0.41 | [-0.52, 1.07] | 0.67 | 0.503 |

**Table S15** Model fit statistics of the multiple linear regression testing the relationship between species richness and the spatial variability of biomass (biomass CV) across spatial scales and different land-use intensities (LUI) in the region Hainich-Dün for the year 2020. All continuous variables were square root transformed, except the variable size. A three-way interaction between species richness, spatial scale and LUI category (low, medium, high) was tested. Interaction terms are indicated with a colon. Adjusted $R^2$ = 0.44, $F_{20,2861}$ = 116.10, p-value <0.001***

| Predictor | Estimate | SE | 95% CI | t-value | p-value |
|---|---|---|---|---|---|
| (Intercept) | 3.09 | 0.69 | [1.74, 4.43] | 4.5 | <0.001*** |
| sqrt(mean_rich) | 0.21 | 0.08 | [0.05, 0.36] | 2.65 | <0.01** |
| scalefield | -1.19 | 0.55 | [-2.27, -0.1] | -2.15 | <0.05* |
| scalefarm | -3.02 | 1.33 | [-5.63, -0.41] | -2.27 | <0.05* |
| LUImedium | -0.46 | 0.75 | [-1.93, 1.01] | -0.62 | 0.538 |
| LUIhigh | -0.58 | 0.71 | [-1.96, 0.81] | -0.82 | 0.413 |
| sqrt(mean_RaoQ) | 0.36 | 0.03 | [0.31, 0.41] | 14.01 | <0.001*** |
| sqrt(mean_TWI) | -0.25 | 0.15 | [-0.54, 0.03] | -1.73 | 0.084 |
| log(size) | 0.26 | 0.02 | [0.23, 0.3] | 14.8 | <0.001*** |
| sqrt(mean_rich):scalefield | 0.23 | 0.09 | [0.05, 0.42] | 2.45 | <0.05* |
| sqrt(mean_rich):scalefarm | 0.65 | 0.24 | [0.17, 1.12] | 2.67 | <0.01** |
| sqrt(mean_rich):LUImedium | 0.09 | 0.13 | [-0.17, 0.34] | 0.65 | 0.514 |
| sqrt(mean_rich):LUIhigh | 0.09 | 0.13 | [-0.16, 0.34] | 0.72 | 0.474 |
| scalefield:LUImedium | -0.38 | 0.87 | [-2.08, 1.32] | -0.44 | 0.658 |
| scalefarm:LUImedium | 1.1 | 1.7 | [-2.23, 4.44] | 0.65 | 0.517 |
| scalefield:LUIhigh | -1.19 | 0.83 | [-2.82, 0.44] | -1.43 | 0.153 |
| scalefarm:LUIhigh | -0.61 | 1.69 | [-3.93, 2.7] | -0.36 | 0.716 |
| sqrt(mean_rich):scalefield:LUI medium | 0.07 | 0.15 | [-0.23, 0.37] | 0.44 | 0.66 |
| sqrt(mean_rich):scalefarm:LUI medium | -0.19 | 0.31 | [-0.8, 0.41] | -0.63 | 0.53 |
| sqrt(mean_rich):scalefield:LUI high | 0.24 | 0.15 | [-0.05, 0.54] | 1.6 | 0.11 |
| sqrt(mean_rich):scalefarm:LUI high | 0.14 | 0.31 | [-0.47, 0.75] | 0.46 | 0.649 |

**Table S16** Model fit statistics of the multiple linear regression testing the relationship between species richness and the spatial variability of biomass (biomass CV) across spatial scales and different land-use intensities (LUI) in the region Hainich-Dün for the year 2021. All continuous variables were square root transformed, except the variable size. A three-way interaction between species richness, spatial scale and LUI category (low, medium, high) was tested. Interaction terms are indicated with a colon. Adjusted $R^2$ = 0.45, $F_{20,2842}$ = 116.30, p-value <0.001***

| **Predictor** | **Estimate** | **SE** | **95% CI** | **t-value** | **p-value** |
|---|---|---|---|---|---|
| (Intercept) | 4.26 | 0.59 | [3.1, 5.43] | 7.18 | <0.001*** |
| sqrt(mean_rich) | 0.12 | 0.07 | [-0.01, 0.26] | 1.81 | 0.071 |
| scalefield | -0.9 | 0.5 | [-1.88, 0.08] | -1.79 | 0.073 |
| scalefarm | -2.03 | 1.12 | [-4.24, 0.17] | -1.81 | 0.07 |
| LUImedium | 0.43 | 0.6 | [-0.74, 1.6] | 0.72 | 0.471 |
| LUIhigh | -1.1 | 0.64 | [-2.37, 0.16] | -1.71 | 0.087 |
| sqrt(mean_RaoQ) | 0.37 | 0.02 | [0.34, 0.41] | 18.95 | <0.001*** |
| sqrt(mean_TWI) | -0.61 | 0.13 | [-0.85, -0.36] | -4.84 | <0.001*** |
| log(size) | 0.22 | 0.01 | [0.19, 0.25] | 14.89 | <0.001*** |
| sqrt(mean_rich):scalefield | 0.18 | 0.08 | [0.02, 0.35] | 2.19 | <0.05* |
| sqrt(mean_rich):scalefarm | 0.39 | 0.19 | [0.01, 0.77] | 2.02 | <0.05* |
| sqrt(mean_rich):LUImedium | -0.1 | 0.1 | [-0.29, 0.1] | -0.95 | 0.341 |
| sqrt(mean_rich):LUIhigh | 0.2 | 0.11 | [-0.02, 0.42] | 1.81 | 0.07 |
| scalefield:LUImedium | 0.38 | 0.72 | [-1.02, 1.79] | 0.53 | 0.594 |
| scalefarm:LUImedium | -0.42 | 1.47 | [-3.31, 2.46] | -0.29 | 0.774 |
| scalefield:LUIhigh | 1.37 | 0.75 | [-0.1, 2.85] | 1.83 | 0.068 |
| scalefarm:LUIhigh | 2.07 | 1.44 | [-0.76, 4.9] | 1.43 | 0.152 |
| sqrt(mean_rich):scalefield:LUI medium | -0.06 | 0.12 | [-0.3, 0.18] | -0.51 | 0.608 |
| sqrt(mean_rich):scalefarm:LUI medium | 0.13 | 0.26 | [-0.37, 0.63] | 0.51 | 0.609 |
| sqrt(mean_rich):scalefield:LUI high | -0.28 | 0.13 | [-0.54, -0.03] | -2.17 | <0.05* |
| sqrt(mean_rich):scalefarm:LUI high | -0.35 | 0.25 | [-0.84, 0.15] | -1.38 | 0.168 |

**Table S17** Effect sizes (eta squared) of multiple linear regressions testing the relationship between species richness and biomass as well as between species richness and the spatial variability of biomass (biomass CV) for all four subsets based on region (SCH: Schorfheide-Chorin, HAI: Hainich-Dün) and year. Effect sizes refer to multiple linear regressions reported in Table S9-16. Effect sizes are bold if above 0.3

| **Parameter** | | **SCH 2020** | **HAI 2020** | **SCH 2021** | **HAI 2021** |
|---|---|---|---|---|---|
| Sqrt(biomass) | **sqrt(mean_rich)** | **0.434853** | **0.496441** | 0.168725 | **0.368693** |
| | scale | 0.003668 | 0.006521 | 0.000642 | 0.007245 |
| | LUI_cat | 0.012935 | 0.022431 | 0.00299 | 0.01895 |
| | sqrt(mean_RaoQ) | 0.073804 | 0.006382 | 0.021277 | 0.009258 |
| | sqrt(mean_TWI) | 0.000875 | 0.013212 | 0.000926 | 0.019605 |
| | log(size) | 0.00025 | 1.55E-05 | 0.000467 | 0.001812 |
| | sqrt(mean_rich):scale | 0.001891 | 0.002269 | 0.000499 | 0.00295 |
| | sqrt(mean_rich):LUI_cat | 0.00375 | 4.71E-05 | 0.003759 | 0.005967 |
| | scale:LUI_cat | 0.006134 | 0.000487 | 0.001684 | 0.001426 |
| | sqrt(mean_rich):scale:LUI_cat | 0.002893 | 0.00146 | 0.004851 | 0.001057 |
| Sqrt(biomass CV) | sqrt(mean_rich) | 0.073453 | 0.064519 | 0.02235 | 0.005212 |
| | **scale** | **0.45285** | **0.378914** | **0.369019** | **0.388417** |
| | LUI_cat | 0.010629 | 0.000822 | 0.00161 | 0.003369 |
| | sqrt(mean_RaoQ) | 0.205283 | 0.023361 | 0.087766 | 0.063973 |
| | sqrt(mean_TWI) | 0.007377 | 0.001464 | 0.000105 | 0.00916 |
| | log(size) | 0.1641 | 0.078435 | 0.064807 | 0.077235 |
| | sqrt(mean_rich):scale | 0.019854 | 0.013986 | 0.003517 | 0.003822 |
| | sqrt(mean_rich):LUI_cat | 0.000431 | 0.005164 | 0.002241 | 0.002375 |
| | scale:LUI_cat | 0.003956 | 0.000568 | 0.001393 | 0.004428 |
| | sqrt(mean_rich):scale:LUI_cat | 0.003684 | 0.001255 | 0.000751 | 0.00251 |

**Table S18:** Model fit statistics of multiple regressions testing BEF relationships based on predicted and measured data in grassland plots of the biodiversity exploratories. Model fit statistics for the response variable predicted biomass (log-transformed): F-Test, $F_{3,196}$ = 61.56, p-value <0.001***, adjusted $R^2$ = 0.48. Model fit statistics for the response variable measured biomass (log-transformed): F-Test, $F_{4,69}$ = 4.271, p-value <0.01**, adjusted $R^2$ = 0.15. Model fit statistics for the response variable predicted biomass CV: F-test, $F_{3,196}$ = 11.13, p-value <0.001***, adjusted $R^2$ = 0.13. Model fit statistics for the response variable measured biomass CV: F-test, $F_{4,69}$ = 0.8502, p-value <0.05, adjusted $R^2$ = -0.01

| Response | Predictor | Estimate | SE | 95% CI | t-value | p-value |
|---|---|---|---|---|---|---|
| Predicted log(biomass) | (Intercept) | 5.79 | 0.09 | [5.62, 5.96] | 66.47 | <0.001*** |
| | richness | -0.04 | 0.00 | [-0.04, -0.03] | -11.01 | <0.001*** |
| | year2021 | 0.21 | 0.05 | [0.11, 0.31] | 4.22 | <0.001*** |
| | exploHAI | 0.10 | 0.07 | [-0.03, 0.23] | 1.52 | 0.13 |
| Measured log(biomass) | (Intercept) | 6.19 | 0.38 | [5.44, 6.94] | 16.45 | <0.001*** |
| | richness | -0.01 | 0.01 | [-0.04, 0.01] | -1.16 | 0.25 |
| | year2021 | 0.40 | 0.19 | [0.02, 0.78] | 2.08 | <0.05* |
| | exploHAI | -0.20 | 0.22 | [-0.96, 0.03] | -0.89 | 0.38 |
| | exploALB | 0.27 | 0.28 | [-0.83, 0.30] | 0.94 | 0.35 |
| Predicted biomass CV | (Intercept) | 5.51 | 1.34 | [2.88, 8.15] | 4.13 | <0.001*** |
| | mean_rich | 0.16 | 0.05 | [0.06, 0.26] | 3.12 | <0.01** |
| | year2021 | -2.01 | 0.77 | [-3.53, -0.49] | -2.61 | <0.01** |
| | exploHAI | 1.25 | 1.00 | [-0.73, 3.23] | 1.25 | 0.21 |
| Measured biomass CV | (Intercept) | 11.19 | 3.30 | [4.60, 17.78] | 3.39 | <0.01** |
| | richness | 0.13 | 0.11 | [-0.10, 0.36] | 1.15 | 0.25 |
| | year | 0.95 | 1.73 | [-2.50, 4.41] | 0.55 | 0.58 |
| | exploHAI | 1.29 | 2.25 | [-3.19, 5.77] | 0.57 | 0.57 |
| | exploALB | 1.95 | 2.57 | [-3.17, 7.08] | 0.76 | 0.45 |

**Table S19** Results of the linear regression which is used as calibration curve for biomass and measurements of the Rising Plate Meter (RPM). Both variables were log-transformed. Model fit statistics: F-test, $F_{1,216}$ = 361.8, p-value <0.001***, adjusted $R^2$ = 0.62. Equation of calibration curve:
$y = 3.11 + 1.06x$

| Response | Predictor | Estimate | SE | 95% CI | t-value | p-value |
|---|---|---|---|---|---|---|
| Log(biomass) | (Intercept) | 3.11 | 0.16 | [2.81, 3.42] | 19.97 | <0.001*** |
| | log(RPM) | 1.06 | 0.06 | [0.95, 1.17] | 19.02 | <0.001*** |
